# Linkers in bitopic agonists shape bias profile among transducers for the dopamine D2 and D3 receptors

**DOI:** 10.1101/2024.01.07.574547

**Authors:** Ana Semeano, Rian Garland, Alessandro Bonifazi, Kuo Hao Lee, John Famiglietti, Wenqi Zhang, Yoon Jae Jo, Francisco O. Battiti, Lei Shi, Amy Hauck Newman, Hideaki Yano

**Author notes:** Corresponding author Hideaki Yano.

## Abstract

Bitopic ligands bind both orthosteric and allosteric or secondary binding sites within the same receptor, often resulting in improvement of receptor selectivity, potency, and efficacy. In particular, for both agonists and antagonists of the dopamine D2 and D3 receptors (D2R and D3R), the primary therapeutic targets for several neurological and neuropsychiatric disorders, bitopic ligand design has proved advantageous in achieving better pharmacological profiles *in vitro*. Although the two pharmacophores within a bitopic ligand are typically considered the main drivers of conformational change for a receptor, the role of the linker that connects the two has not yet been systematically studied for its relevance in receptor activity profiles.

Here, we present a comprehensive analysis of sumanirole and PF592,379-based indole-containing bitopic compounds in agonist activity at D2R and D3R, with a focus on linker chemical space and stereochemistry achieved through testing seven distinct chirally resolved linkers. The current study examines the structure activity relationships (SAR) of these linkers extensively, beyond the conventional level, by characterizing activation of all putative transducers over a 44 min time course. Our multiparametric analysis provides previously unappreciated clarity of linker-dependent effects, highlighting the utility of this applied comprehensive approach and the significance of linker type in the shaping of transducer bias profiles.

## Introduction

Agonists of dopamine D2 and D3 receptors (D2R and D3R) are used in the treatment of Parkinson’s Disease (PD) and other motor disorders. Despite their differential expression in the brain, both D2R and D3R engage in molecular signaling through the same Gi/o proteins, inhibiting adenylyl cyclase-mediated cAMP production and activating G protein-gated inwardly rectifying potassium (GIRK) channels [1, 2]. Homing in on agonist specificity for biased targeting (i.e., preferential activation of one signaling pathway over another) of a receptor and its associated signal transducer subtypes [3], particularly by agonists with higher potency, would advance the field by affording a greater understanding of the nuances of D2R and D3R activation. The identification of such specific signaling targets has powerful potential to yield cell-type specific signaling activities. However, the nearly identical orthosteric binding site shared by the two receptors, in combination with high sequence homology between D2R and D3R transmembrane domains, presents a significant challenge in efforts to design highly selective therapeutics [4–6].

Our previous research has described the development of the D2-preferential agonist sumanirole (SM) and associated selective bitopic compounds in D2R [7–9], and of the D3R-preferential agonist PF592,379 (PF) and associated selective bitopic ligands in D3R [8, 10, 11]. Through previous work, we have identified ideal linker length, chemical composition and substitutions, with consequent stereochemistry, and optimal secondary pharmacophores for these novel bitopic agonists [8, 11–15]. Beyond this, we have revealed the potential for use of biased pharmacology through comparison of G-protein versus β-arrestin signaling [7, 9, 11], and Gi1 versus GoA signaling [7, 9]. These studies extend ligand-biased signaling beyond β-arrestin-G protein selectivity to functional selectivity among G proteins. While much effort has gone into thorough characterization of transducers in recent years [3, 16, 17], there is a paucity of focus on Gi subtype bias in D2R and D3R, which warrants further investigation. As there are certain Gi subtypes which are expressed more highly in the brain (e.g., via GoA, GoB, Gz activation; [18]), subtype-biased pharmacology offers the potential for development of more centrally oriented targeting.

Within bitopic ligand structure, linker composition is often overlooked for its relevance in transducer activation, particularly in functional selectivity. Given the different pharmacological effects associated with linkers indicated by our previous work [8, 10, 11, 13], as well as work by other groups [19–21], we took this idea further by introducing a variety of chiral aliphatic cyclic linkers. Because the secondary pharmacophore has previously been implicated as having a critical role in signaling bias [22], our current study puts special emphasis on the linker, which not only serves to orient the secondary pharmacophore, but also itself interacts extensively with the receptor to influence its conformation, modulating subtype selectivity and functional profiles [10, 23]. Through this resolution of chiral linkers, which also includes comparable diastereomeric pairs, we enhance our understanding of spatial arrangements of the linker and interacting vestibule between the orthosteric and allosteric sites in both D2R and D3R.

In this study, we delve further into the intricate interplay between ligand structure and the activity of transducers subtypes in the context of GPCRs. We achieve this by conducting comprehensive functional characterization of both PF and SM ligand series in D2R and D3R, assessing their activation profiles and kinetics across six G-protein subtypes - GoA, GoB, Gz, Gi1, Gi2, and Gi3 – and recruitment of βarr1 and βarr2. Our primary goal is to study the selectivity and affinity of these ligands, ultimately uncovering unique transducer activation profiles within both D2R and D3R, and guide future ligands optimization campaigns.

## Methods

### Plasmids

Transfectant plasmids include human dopamine D2 (D2R) and D3 (D3R) receptors, either untagged or fused with Renilla luciferase 8 (RLuc8); G protein subunits: GαoA, GαoB, Gαz, Gαi1, Gαi2, and Gαi3 fused with RLuc8 and Gγ2 fused with yellow fluorescent protein variant Venus. Unlabeled β1 subunit was also co-transfected in all experiments. Venus fused β-arrestin 1 or β-arrestin 2 with RLuc8 fused D2R or D3R were used to monitor β-arrestin recruitment. GRK2 was co-transfected to promote enhanced phosphorylation necessary for the β-arrestin recruitment.

### Transfection

Human embryonic kidney cells 293T (HEK 293T) were seeded at 3 million cells per 10-cm plate and cultured in Dulbecco’s modified Eagle’s medium (DMEM) with 10% fetal bovine serum (FBS), 1% penicillin-streptomycin, and 2 mM L-glutamine at 37 °C and 5% CO_2_ 95 % moisturized air. HEK293T cells were transiently transfected with a total of 15 ug of plasmid cDNA of each above mentioned construct using polyethyleneimine (PEI) at a ratio of 2:1 (PEI:total DNA by weight) with an incubation time of ∼48 hours.

### Bioluminescence Resonance Energy Transfer (BRET) Studies

The BRET-based Go protein activation and β-arrestin recruitment assays were performed as described previously [9, 24]. Go protein activation assay uses Renilla luciferase 8-fused Gα_oA_ and mVenus-fused Gγ_2_ as the BRET pair. β-arrestin recruitment assay uses RLuc8-fused D_3_R or D_2_R and mVenus-fused β-arrestin2 as the BRET pair. After ∼48 h of transfection, cells were washed, harvested, and resuspended in 1X Phosphate-buffered saline (PBS) + 0.1% glucose + 200 µM sodium bisulfite (NaBi) antioxidant and anti-browning reagent. Cells were plated in 96-well plates (White Lumitrac 200, Greiner bio-one, Monroe,NC, USA) at a density of approximately 200,000 cells per well. BRET luciferase substrate coelenterazine H (5 uM) was then added at a rate of 1 well per second. Reference D_2_/D_3_ agonist dopamine (Tocris Bioscience, Minneapolis, MN, USA), test ligands, and vehicle controls were manually added 3 min later via multichannel pipette. BRET signal was measured using a Pherastar FSX plate reader (BMG Labtech, Cary, NC, USA). For kinetic experiments, cells were incubated at 37 °C within the Pherastar FSX plate reader, with BRET signal measurements taken every 2 min during 44 min.

### Molecular Docking

The molecular modeling and docking were carried out with the Schrodinger Suite (version 2022-1). The pKa predictions of the protonation state of the ligands were performed using Epik [25]. The 3D structures of the ligands were then generated using LigPrep. The active structure of dopamine D3 receptor (D3R) was taken from the crystal structure of the receptor-G protein complexes (PDB ID 7CMV, [26]), and the protein preparation protocol was performed to prepare the D3R model. Finally, the induced fitted docking (IFD) protocol [27] was used to perform molecular docking. The final binding poses were selected based on the best IFD scores obtained.

### D2R and D3R Bitopic Ligands

All bitopic ligands consist of a primary pharmacophore (PP), either a Sumanirole (SM, D2R-preferential full agonist PP for SM series compounds), or PF592,379 (either PF*_R,S_*or PF*_S,S_* D3R-preferential full agonist PP, with PF*_S,S_* for PF bitopic series compounds), attached to an indole-2-amide moiety secondary pharmacophore (SP) via one of the unique linkers as described below.

Bitopic Ligands: SM-series bitopic ligands include VK01-140 (SM-A), with an alkyl linker (Linker A); AB08-41-P1 (SM-B*_cis_*) and AB08-41-P2 (SM-B*_trans_*), with a chiral cyclopentyl linker (Linker B); cis-AB10-90 (SM-C*_cis_*) and trans-AB10-89 (SM-C*_trans_*), with a chiral cyclohexyl linker (Linker C); AB05-98A (SM-D*_R,S_*) and AB05-98B (SM-D*_S,R_*) with a chiral cyclopropyl linker (Linker D). PF-series bitopic ligands include AB04-87 (PF*_R,S_*-A), with trans-PF PP and an alkyl linker (Linker A); AB04-88 (PF*_S,S_*-A) with cis-PF PP and an alkyl linker (Linker A); FOB04-04A (PF*_S,S_*-B*_cis_*) and FOB04-04B (PF*_S,S_*-B*_trans_*), with a chiral cyclopentyl linker (Linker B); AB08-57A (PF*_S,S_*-C*_cis_*) and AB08-57B (PF*_S,S_*-C*_trans_*), with a chiral cyclohexyl linker (Linker C); FOB02-04B (PF*_S,S_*-D*_R,S_*) and FOB02-04A (PF*_S,S_*-D*_S,R_*), with a chiral cyclopropyl linker (Linker D). Synthesis, resolution and characterization of PF*_S,S_*, PF*_R,S_*, SM-C*_cis_* and SM-C*_trans_* are described in the S.I., and all the other compounds have been previously reported [8].

### Data Analysis

BRET ratio was calculated as the ratio of Venus (530 nm) over RLuc8 (480 nm) emission. BRET signals were detected every 2 min during a 44 min period using the Pherastar FSX plate reader incubated at 37°C. A script (Python) was developed for data processing, used to transform and organize kinetic data prior to regression analysis. Dose-response curves were generated using a non-linear fit log(agonist) vs. BRET ratio response using Prism 10 (GraphPad Software, San Diego, CA, USA) and presented as mean ± SEM. Data were collected from at least 4 independent experiments per condition. Detection of outliers when fitting data with nonlinear regression was performed using ROUT method, combining robust regression and outlier removal, with Q=1% outliers exclusion and fitting method of least squares regression. All data was transformed for individual BRET ratio values and further normalized to the maximal response by dopamine at 10 min as 100% and the minimal response by dopamine at 10 min as 0%, within each respective transducer. E_max_ and pEC_50_ parameters were directly obtained from Non-linear fit of normalized of transformed data and multicomparison data analyses at the 10 min and 40 min time point were conducted. To evaluate whether ligands exhibited G protein subunit signaling bias, bias factors were calculated as reported previously [28]. This method yields bias factors similar to the operational model [9]. Briefly, 1) Δ log(*E_max_*/*EC*_50_) value for each transducer was calculated by subtracting the log(*E_max_*/*EC*_50_) value of the agonist by that of the reference ligand (dopamine).

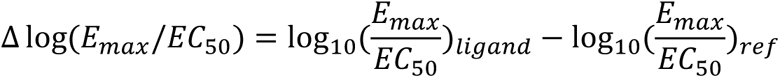

2) ΔΔ log(*E*_*max*_/*EC*_50_) consists of the subtraction of Δ log(*E_max_*/*EC*_50_) between two transducers. The comparison between the two transducers shows the bias direction of a certain ligand when activating these two transducers.

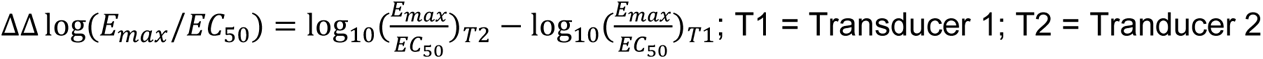

### Statistical Analysis

Triplicate of at least four independent experiments were performed per each condition. Statistical grouped analyses on E_max_ and pEC_50_ were conducted using two-way ANOVA with Tukey’s multiple comparisons within each timepoint, comparing across all different agonists, with 95% confidence interval and statistically significant P <0.0332(*), <0.0021(**), <0.0002(***), <0.0001(****).

## Results/Discussion

### 1. Enhancement of potency through bitopic strategy

We begin with functional characterization of PF592,379 (PF) and sumanirole (SM), the two primary pharmacophores in a series of bitopic ligands used in this study. PF can be resolved into two diastereomers at the 2 position of the morpholine ring, namely PF*_S,S_* and PF*_R,S_*. Using these scaffolds, we have designed bitopic ligands with an indole motif secondary pharmacophore tethered to the primary pharmacophore by a 4-carbon chain linker structure, termed linker A. This approach offers insight into how the bitopic nature of ligands can contribute to modulating the pharmacological profile of primary pharamacophores in both SM and PF bitopic ligand series, whose structural design is shown in Figure 1.

**Figure 1.**
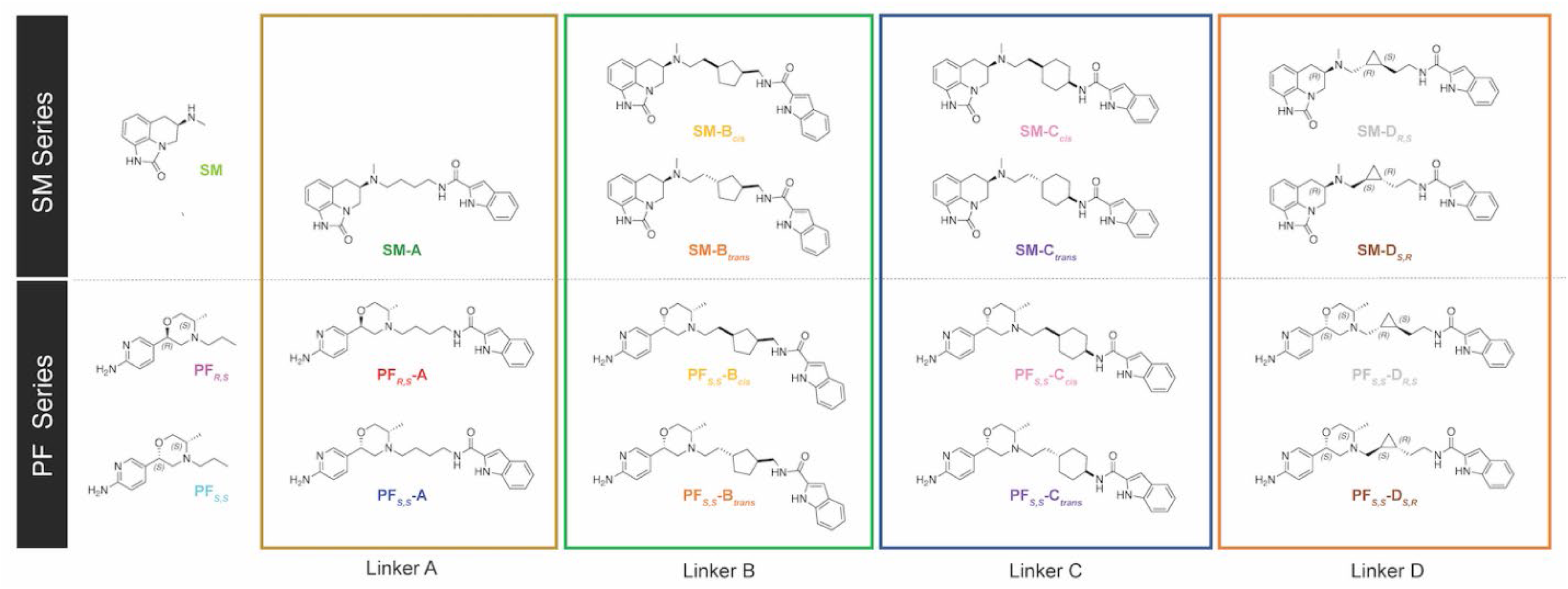
Drug design for sumanirole and PF592,379 series of bitopic ligands. Bitopic ligand series based on primary pharmacophore sumanirole or PF592,379 moiety was synthesized using different chiral linkers for indole attachment: butyl (Linker A), cyclopentyl (Linker B), cyclohexyl (Linker C), and cyclopropyl (Linker D).

Assessment of potency (i.e., pEC_50_) of the D2R preferential agonist SM, reveals a range of 6.2-7.3 in D2R (Table 1) and 5.5-7.1 in D3R (Table 2) across various transducers, with no significant potency difference (less than 5-fold) between D2R and D3R. SM showed comparable potency to the endogenous agonist dopamine (DA) in D2R. However, in D3R, SM had lower potency than DA, with a decrease in G-protein activation up to 45-fold, and a remarkable decrease between 299-782-fold in β-arrestin recruitment. Introduction of the indole secondary pharmacophore to SM via linker A, designated as ligand SM-A, results in a identical potency profile, with less than 10-fold potency difference observed between SM-A and SM in both D2R (Supplementary Figure S1, A-H) and D3R (Supplementary Figure S1, I-P) across various transducers, with the exception of D3R-βarr2 (>10-fold). In contrast to our previously reported findings [7], the data for SM-A presented in this study reveal distinct pharmacological outcomes in GoA, in particular a 58-fold reduction in potency for D2R-GoA signaling.

**Table 1.**
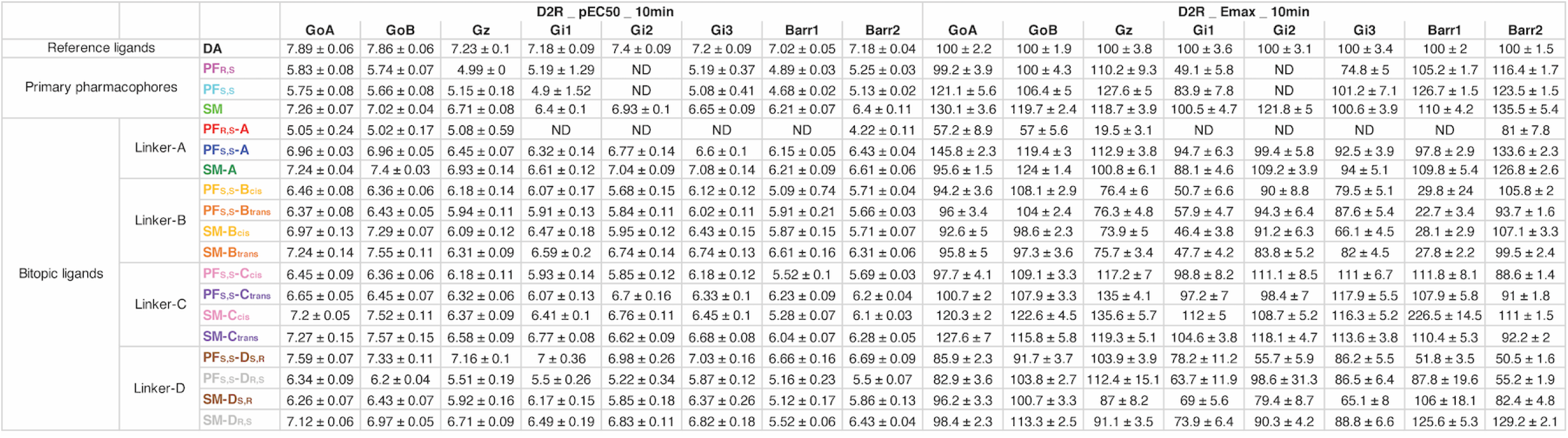
– Potency and efficacy of bitopic sumanirole and PF592,379 ligands for D2R activities at 10 min.

**Table 2.**
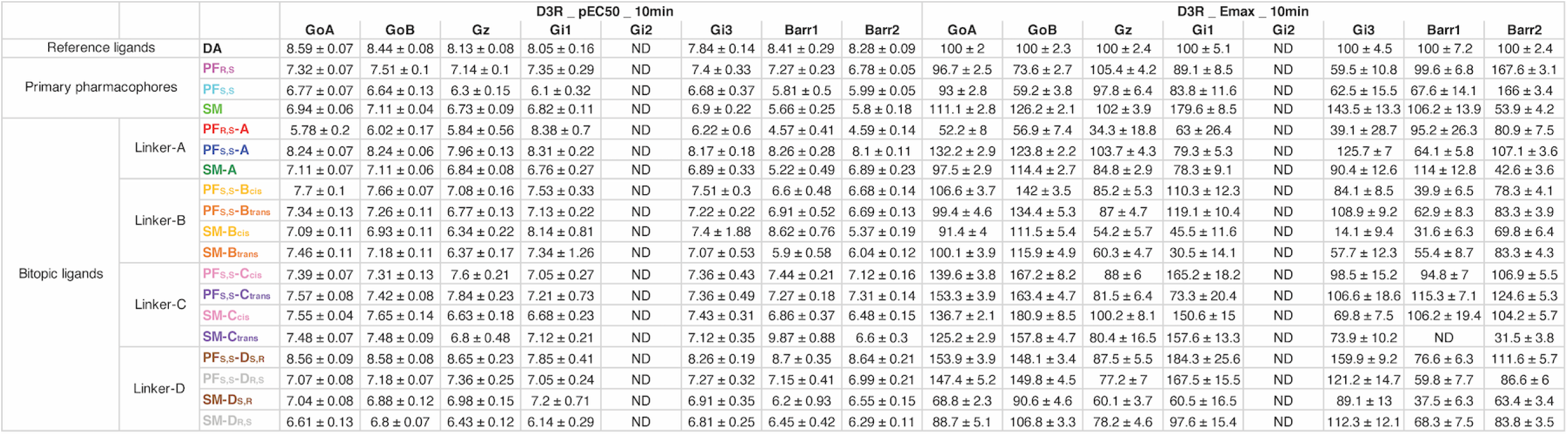
– Potency and efficacy of bitopic sumanirole and PF592,379 ligands for D3R activities at 10 min.

In terms of efficacy for SM, the parent ligand exhibited up to 35% higher E_max_ relative to DA in D2R and E_max_ up to 80% higher in D3R. The bitopic ligand SM-A showed slightly lower efficacy than SM, with a difference of 12-34% in D2R and 11-100% in D3R.

When analyzing potency of PF diastereomers, both PF*_S,S_* and PF*_R,S_* were more potent activators of D3R than D2R. Across all G-proteins, potency of PF*_S,S_* ranged from 7-40 times higher, while potency of PF*_R,S_* ranged from 31-240 times higher in D3R over D2R. These findings align with prior reports establishing selectivity of a diastereomeric mixture of PF, and its PF*_R,S_* isomer for D3R over D2R [10, 11, 29]. Furthermore, PF*_R,S_* is slightly more potent than PF*_S,S_* across all transducers in both D2R and D3R. For example, in GoA activation, PF*_S,S_* and PF*_R,S_* showed pEC_50_ of 5.75 and 5.83 for D2R (Table 1) and pEC_50_ of 6.77 and 7.32 for D3R (Table 2), respectively. PF*_S,S_* and PF*_R,S_* exhibited potency significantly lower than DA, by 19-386-fold in both D2R and D3R across all transducers at the 10 min mark, the timepoint of analyzed results in section 1-5. We further investigated the behavior of PF diastereomers as primary pharmacophores within a bitopic ligand. A substantial potency shift resulted from the introduction of an indole secondary pharmacophore via linker A attachment to the PF primary pharmacophores. The bitopic ligand PF*_S,S_*-A exhibited a 16-34-fold increase in potency in D2R (Supplementary Figure S2, A-H) and a remarkable 31-163-fold increase in potency in D3R (Figure 2, A-H), across different transducers (excluding Gi2) as compared to the parent ligand PF*_S,S_*. The observation of a large increase in potency for PF*_S,S_*-A is consistent with a previous report on activation of GoA and β-arrestin by the same ligand (referred to as AB04-88) [11]. In contrast to the observed effects of SM ligands, the pronouced potency shift between PF*_S,S_*-A and PF*_S,S_* can be attributed to inherent disparities between the SM-scaffold and the PF-scaffold. Interestingly, the same chemical modification applied to the PF*_R,S_* diastereomer resulted in decreased potency of its respective bitopic ligand PF*_R,S_*-A; a <10-fold decrease in D2R and a 15-35-fold decrease in D3R compared to PF*_R,S_*. Where pEC_50_ or E_max_ parameters could not be determined (indicated as ND) as the result of an inability to fit regression analyses to low potency shift curves, or due to the absence of a robust BRET signal (as observed in D3R-Gi2 and D3R-βarr1), cases are marked explicitly in Supplementary Table 2. Finally, addition of the secondary pharmacophore to the PF diastereomers resulted in a drastic separation of potency between the bitopic diastereomeric pair PF*_S,S_*-A and PF*_R,S_*-A across different transducers. Specifically, for GoA, GoB, and Gz, a 24-88-fold potency separation in D2R and 131-291-fold separation in D3R was observed. This represents a remarkable difference when compared to the primary pharmacophores PF*_S,S_* and PF*_R,S_*, whose potency difference does not exceed 7-fold for either D2R or D3R.

**Figure 2.**
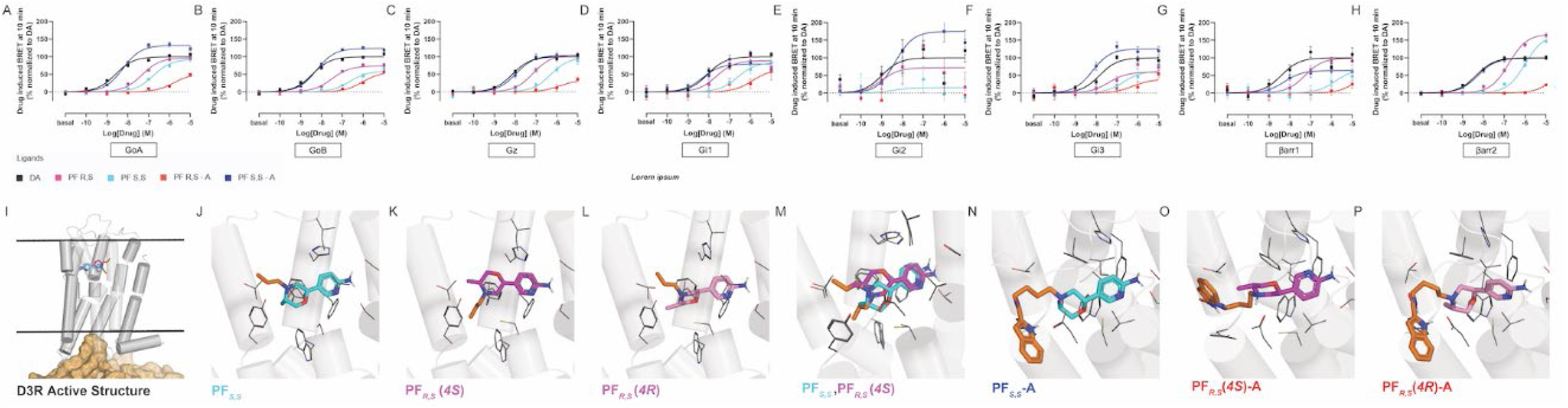
D3R-transducer activation profile of bitopic PF592,379 ligands with butyl linker at 10 minutes and their docking within the binding pocket. Concentration-response curves of activation BRET between Gα-subtype-RLuc and Gγ-Venus (A-F), and D3R-RLuc and β-arrestin-subtype-Venus (G-H). Curves are presented as a percentage of the maximal response of DA with means ± SEM (n ≥ 4). Docking interactions within the binding pocket (I-P). (2R, 4S, 5S)-PF592,379 (magenta) and (2S, 4R, 5S)-PF592,379 (cyan) at the orthosteric binding site (OBS) of D3R-Gi1 (I). Propyl chain of (2S, 4R, 5S)-PF592,379 pointing toward the extracellular side (J,M). For (2R, 4S, 5S)-PF592,379, the same chain pointing to the cavity between TM6 and TM7 (K,M). In two configurations (4S vs. 4R) of trans-PF (K,L), the favorable hydrophobic contact observed for (2R, 4S, 5S)-PF592,379 (K). Docking poses of bitopic compounds (N-P) for AB04-88 (N) and AB04-87 (O-P) with favorable interactions observed for AB04-88 (N).

Regarding the efficacy of PF diastereomers relative to DA, PF*_S,S_* and PF*_R,S_* showed E_max_ up to 135% in D2R (except for Gi1 and Gi3, with E_max_ below 84%) (Supplementary Figure S2, A-H), but as low as 40% in D3R (except for βarr2 with E_max_ 167%) (Figure 2, A-H). When PF diastereomers and bitopic ligands were compared, PF*_S,S_* and PF*_S,S_*-A exhibited similar efficacy in several transducers. However, in D2R-Gi2 and D2R-βarr1, there was 29-36% lower efficacy for PF*_S,S_*-A compared to PF*_S,S_* (Table 1), while in D3R-GoA, D3R-GoB, and D3R-Gi3 activation, PF*_S,S_*-A showed 39-65% higher efficacy compared to PF*_S,S_* (Table 2). PF*_R,S_*-A showed 35-111% lower efficacy across all transducers in D2R and 45-87% lower efficacy in D3R-GoA, D3R-Gz, and D3R-βarr2 compared to its parent ligand PF*_R,S_*. Finally, the addition of the secondary pharmacophore to the PF diastereomers led to improved efficacy in PF*_S,S_*-A relative to its diastereomer PF*_R,S_*-A. Notably, a remarkable 62-93% increase in E_max_ for PF*_S,S_*-A over PF*_R,S_*-A was observed in D2R, with a similar increase of 67-87% in D3R across Gi subtypes (except for Gi1). PF*_S,S_* showed a more modest difference, between 20-35% higher E_max_ over PF*_R,S_* in D2R (except for GoB and βarr2, E_max_ difference <10%), with no significant E_max_ difference observed in D3R across all transducers.

To support these results, ligand docking simulations at the active structure of D3R (PDB 7CMV) were performed using the induced-fit docking with enhanced sampling protocol from Schrodinger Suite 2022-1. The docking process accounts for both protonation states of the charged nitrogen in the morpholinium moiety (Figure 2 I-P). Pharmacological results reveal that both PF*_R,S_* and PF*_S,S_* exhibit binding affinity for the orthosteric binding site (OBS) of the D3R-Gi1 complex (Figure 2 I-P). Analysis of molecular docking poses indicate that the propyl chains of PF*_R,S_* extend towards the cavity between transmembrane helix 6 (TM6) and transmembrane helix 7 (TM7), while the same chains of PF*_S,S_* point towards the extracellular side (Figure 2 I-P). Furthermore, when considering specific isomers, a favorable hydrophobic contact is observed exclusively for PF*_R,S_* (Figure 2 I-P). Additionally, the docking poses of bitopic ligands, namely PF*_S,S_*-A and PF*_R,S_*-A, are depicted with favorable interactions apparent between PF*_S,S_*-A and surrounding residues (Figure 2 I-P). These findings are consistent with the enhanced potency and efficacy of PF*_S,S_*-A, providing important insight into binding characteristics and their potential pharmacological implications for the investigated ligands at the D3R-Gi1 complex. Based on these results, PF*_S,S_*-A emerged as the more promising bitopic ligand among the two diastereomers, and was selected as the reference scaffold for assessment of the performance by ligands with further linker modifications.

In summary, it is evident that PF diastereomers exhibit higher potency in D3R compared to D2R, with no significant tendency for greater potency of PF*_R,S_* over PF*_S,S_*. Secondary pharmacophore addition yielded a dramatically different trend, with a shift in potency that increased for PF*_S,S_*-A and decreased for PF*_R,S_*-A. PF*_S,S_*-A was determined to be more potent than both its parent ligand PF*_S,S_* and its diastereomer PF*_R,S_*-A, a relationship consistent in both D2R and D3R across all transducers, due to favorable interactions with surrounding residues in the OBS. This resulted in an amplified separation of potency between these bitopic PF diastereomers, which is up to 80 times greater than the separation observed between the two parent ligands. Moreover, the biotopic strategy employed for PF*_S,S_*-A enhanced its selectivity for D3R agonist activity. PF*_S,S_*-A is also more efficacious than PF*_R,S_*-A, with PF*_S,S_*-A demonstrating at least 67-110% higher E_max_ of G-protein activation in both D2R and D3R compared to PF*_R,S_*-A, while PF*_S,S_* shows no more than 35% higher E_max_ when compared to PF*_R,S_*. The use of a biotopic strategy did not lead to any significant improvement in SM potency in either D2R or D3R, and no significant difference. was observed when compared to SM-A.

### 2. Impact of the linker on potency and efficacy within SM ligand series

We next investigated the pharmacological effects of linker type in the SM series of bitopic ligands. Besides the simple butyl linker (referred to as Linker A), we tethered the indole secondary pharmacophore to both SM and PF scaffolds using six distinct aliphatic linkers: 1,3-disubstituted cyclopentyl (referred to as Linker B), 1,4-disubstituted cyclohexyl (referred to as Linker C), and *trans*-cyclopropyl (referred to as Linker D) (Figure 1). In the case of cylopentyl and cyclohexyl linkers, both relative *cis*- and *trans*- distereosiomers have been characterized, for the cyclopropyl linker the optimal *trans*-stereochemistry has been used, and both its *trans*- isomers have been resolved [8] and tested. For the SM series of bitopic ligands discussed in this section, analyses were made with SM-A as the reference ligand.

In the potency comparison, SM-A demonstrated equivalent potency in D2R and D3R, with pEC_50_ ranging between 6.2-7.3 in D2R (Table 1), and 5.8-7.1 in D3R (Table 2), across different transducers. SM-A was as potent as the reference agonist dopamine (DA) in D2R (Figure 3, A-H), but 20-30-fold less potent than DA in D3R (except for βarr1) (Supplementary Figure S3, A-H), Bitopic SM ligands containing linkers B, C and D (i.e., SM-B*_cis_*, SM-B*_trans_*, SM-C*_cis_*, SM-C*_trans_*, SM-D*_S,R_*, and SM-D*_R,S_*) showed no significant difference in potency compared to SM-A for most transducers in D2R and D3R. However, in D3R-βarr1 recruitment by SM-B*_cis_*, SM-C*_trans_*, SM-D*_S,R_*, and SM-D*_R,S_*, a significant potency difference of at least 200-fold higher than that of SM-A was observed. Evaluating transducer profiles, βarr1 potency was the most positively impacted by SM linker modifications. Furthermore, bitopic SM ligands consistently activate Go subtypes (i.e., GoA and GoB) with pEC_50_ values higher than 7 in both receptors (except SM-D*_S,R_* in D2R and SM-D*_R,S_* in D3R).

**Figure 3.**
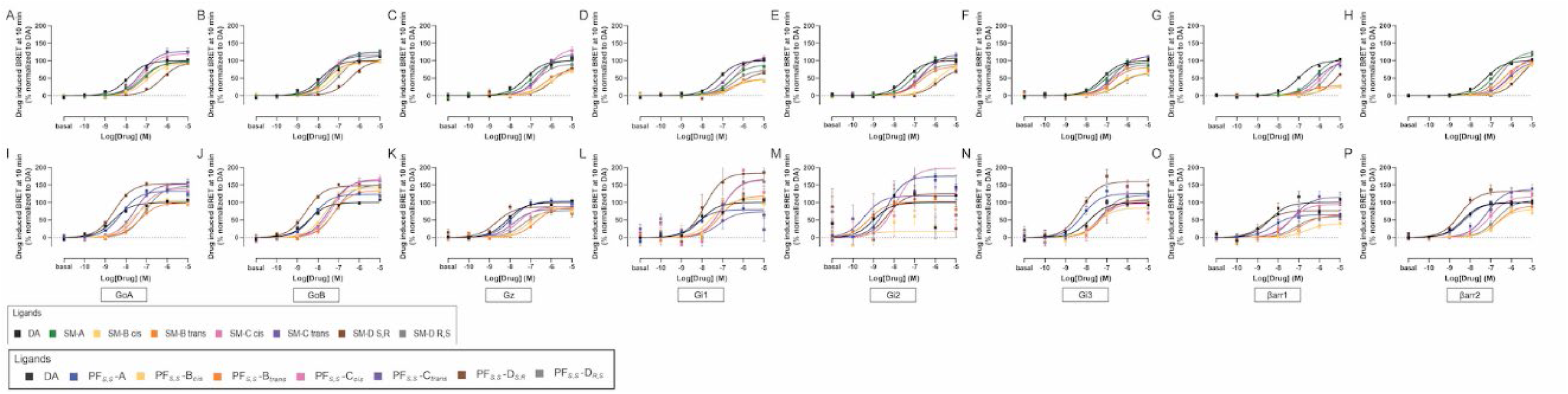
D2R- and D3R-transducer activation profile of bitopic sumanirole and PF592,379 ligands with various linkers at 10 minutes. Concentration-response curves for bitopic SM-ligands in D2R Gα-subtype activation (A-F), and β-arrestin-subtype recruitment (G-H). Concentration-response curves for bitopic PF-ligands in D3R Gα-subtype activation (I-N), and β-arrestin-subtype recruitment (O-P). Curves are presented as a percentage of the maximal response of DA with means ± SEM (n ≥ 4).

In the efficacy comparison, SM-A showed a profile similar to that of DA in D2R (Table 1), with slightly lower E_max_ (less 30% reduction) compared to DA in D3R (Table 2) across most transducers. However, SM-A showed significantly lower efficacy than DA in activation of D3R- βarr2 (E_max_=43%). Compared to other linkers in D2R, linker C ligands (i.e., SM-C*_cis_* and SM-C*_trans_*) showed significantly higher efficacy than SM-A in most transducers, with an E_max_ increase of more than 30% in D2R-GoA, -Gz and -βarr1. Conversely, linkers B and D led to a decrease of E_max_ compared with linker A across all transducers, with the most pronounced difference observed between linker A and linker B ligands. Specifically, SM-B*_cis_* and SM-B*_trans_* demonstrated weaker activation of D2R-βarr1 (82% E_max_ reduction) and D2R-Gi1 (40-42% E_max_ reduction) compared to SM-A. For other transducers and ligands, the E_max_ difference was less than 30%. In D3R, the same pattern of stronger activation by linker C ligands and weaker activation by linker B and D ligands relative to SM-A, was also observed across various transducers. Notably, SM-C*_cis_* exhibited 67-73% higher E_max_ in D3R-GoB, -Gi1, -βarr1 and -βarr2 compared to SM-A, and both SM-B*_cis_* and SM-B*_trans_* exhibited 21-76% lower E_max_ in D3R-Gz, - Gi1, Gi3 and -βarr1 compared to SM-A.

Assessing transducer profiles, in D2R, βarr2 consistently exhibited lower E_max_ with Linkers B, C, and D compared to Linker A. In contrast, βarr2 consistently showed higher E_max_ with Linkers B, C, and D compared to Linker A in D3R, exemplar of a prominent discrepancy in response to bitopic SM ligands by D2R and D3R.

In summary, among the bitopic SM ligands studied here, those containing linker C (i.e., SM-C*_cis_* and SM-C*_trans_*) consistently exhibited enhanced potency compared to SM-A across all transducers in D3R. SM-C*_cis_* and SM-C*_trans_* also demonstrated the highest efficacy across various transducers in D2R and D3R. In contrast, other bitopic SM ligands containing linkers B and D showed lower efficacy across all G-proteins in both D2R and D3R. Moreover, with respect to β-arrestin recruitment, linkers B and D improved efficacy for D3R-βarr2, but not for D2R-βarr2.

### 3. Impact of the linker on potency and efficacy within PF ligand series

With a similar approach to that described in the previous section, we investigated the impact of linker type within the chemical structures of ligands belonging to the PF series through comparison of PF*_S,S_*-A with other bitopic PF-derivative ligands. In the potency comparison, PF*_S,S_*-A showed D3R selectivity, with 19-69-fold potency over D2R across all transducers. The potency of PF*_S,S_*-A is comparable to DA in D3R (Figure 3, I-P) and was approximately 10-fold less than DA in D2R (Supplementary Figure S3, I-P). In D2R (Table 1), PF*_S,S_*-A as well as the majority of bitopic PF ligands showed pEC_50_ <7 across all transducers. PF*_S,S_*-D*_S,R_*, however, had pEC_50_ >7, making it the most potent for D2R among bitopic PF ligands. Further evaluation of transducer profile indicated that linker structure had greater impact in D2R-Gi2; most bitopic ligands showed a significant potency difference as compared to PF*_S,S_*-A, with ΔpEC_50_≥1 at 10 min. In D3R (Table 2), despite the vast majority of ligands having pEC_50_ between 7 and 8, the potency of PF*_S,S_*-D*_S,R_* was still higher, with a pEC_50_ greater than 8 across all transducers (except Gi1), and comparable to that of PF*_S,S_*-A. On the other hand, PF*_S,S_*-D*_R,S_* exhibited the lowest potency, lower than that of PF*_S,S_*-A in both D2R and D3R by 4 to 36-fold, at 10 min. For ligands with other linkers (i.e., linkers B and C) there was no significant potency difference compared to PF*_S,S_*-A across transducers for either D2R or D3R.

In the efficacy comparison, for D2R (Table 1), PF*_S,S_*-A was significantly higher than DA with 46 and 34% greater activation of GoA and βarr2, respectively. When compared to other linkers, we observed that linker C diastereomers (i.e., PF*_S,S_*-C*_cis_* and PF*_S,S_*-C*_trans_*) had an E_max_ up to 25% higher than PF*_S,S_*-A in most transducers (i.e. Gi1, Gi2, Gi3, Gz, and βarr1). Conversely, linkers B and D led to a decrease in E_max_ across all transducers, up to 63% lower in G-proteins and 83% lower in β-arrestins compared to linker A (i.e., PF*_S,S_*-A). For D3R (Table 2), PF*_S,S_*-A demonstrates 20-30% higher efficacy compared to DA in D3R-GoA, -GoB, and -Gi3, but 20-30% lower efficacy in D3R-Gi1 and -βarr1. When compared with other linkers, linkers C and D yielded the highest maximal response for most G protein subtypes. More specifically, PF*_S,S_*-C*_cis_*, PF*_S,S_*-D*_S,R_*, PF*_S,S_*-D*_R,S_* showed a significant enhancement in E_max_, between 86-105% for D3R-Gi1 and between 15-43% for D3R-GoA and -GoB compared to PF*_S,S_*-A. The low E_max_ of PF*_S,S_*-A observed for the D3R-Gi1 transducer resulted in occurance of the greatest efficacy differences between PF*_S,S_*-A and other bitopic PF ligands.

Further evaluation of transducer profiles suggested stronger Gz activation by PF*_S,S_*-C*_cis_*, PF*_S,S_*- C*_trans_*, PF*_S,S_*-D*_S,R_* and PF*_S,S_*-D*_R,S_* in D2R as compared to D3R, in which E_max_ of the ligands ranged between 104-135% in D2R-Gz. In contrast, D3R-Gz activation showed an E_max_ of 77-88%. Conversely, GoA, GoB, and Gi1 activation by the same four ligands was favored in D3R over D2R, with E_max_ between 140-184% in D3R compared to 64-109% in D2R. Furthermore, D3R-GoA and -GoB activation led to high E_max_ (99-181%) with low variability among all bitopic PF ligands. In terms of β-arrestin recruitment, βarr1 was the most negatively impacted by the linker modification (i.e. linker B and D) in both D2R and D3R, a reversed effect of that described for the SM series in the previous section. Specifically, a decrease in E_max_ between 23-50% in PF*_S,S_*-B*_cis_* and PF*_S,S_*-B*_trans_* and between 52-88% in PF*_S,S_*-D*_S,R_* and PF*_S,S_*-D*_R,S_* was observed for D2R and D3R-βarr1. Interestingly, ligands with the linker C modification, PF*_S,S_*-C*_cis_* and PF*_S,S_*- C*_trans,_* did not appreciably impact D2R or D3R-βarr1 activation.

Our findings indicate that linker structure has significant influence on potency and efficacy across various receptors and transducers. Of the bitopic PF ligands investigated, only PF*_S,S_*-D*_S,R_* demonstrated improved potency compared to PF*_S,S_*-A, consistent in both D2R and D3R. PF*_S,S_*- D*_S,R_* was also the ligand with the highest efficacy in D3R-Gi1, although other bitopic ligands also demonstrated improved efficacy as compared to PF*_S,S_*-A, particularly in activating D3R-Gi1, GoB and βarr1. Furthermore, linkers C and D in bitopic PF ligands favored the activation of D3R-Gi1 and D3R-GoB over D2R, while also favoring D2R-Gz activation over D3R. Some of these findings align with those observed in the SM series, indicating the similar role of linker-C-indole moiety in conferring higher efficacy, particularly in activating D3R-Gi1, GoB and βarr1. Further comparisons between the SM and PF series will be discussed in the subsequent section.

### 4. Pharmacological comparison between SM and PF ligand series

Next we compared bitopic ligands containing two different primary pharmacophores (i.e., SM and PF) with matched linkers (i.e., PF*_S,S_*-A vs. SM-A, PF*_S,S_*-B*_cis_* vs. SM-B*_cis_*, etc.) Through a series of pairwise comparisons, we analyzed nuanced effects of the linker-indole moiety on a primary pharmacophore, and compared how these effects differ between SM- and PF-series compounds.

With respect to potency differences, comparison of parent ligands in D2R (Table 1) revealed that PF*_S,S_* potency was significantly lower (18-37-fold) than that of SM across all transducers (except for Gi2), whereas in D3R (Table 2), no significant potency difference was observed between PF*_S,S_* and SM. Conversely, when bitopic linker-A ligands were compared in D2R, PF*_S,S_*-A and SM-A showed no potency difference; whereas in D3R, PF*_S,S_*-A showed 13-142-fold greater potency than SM-A. When evaluating the other bitopic PF vs. SM pairs of identical linkers in D2R, bitopic PF ligands showed lower non-significant potency compared to their specific linker pair from SM series (except for PF*_S,S_*-D*_S,R_ vs.* SM-D*_S,R_*). In contrast, evaluation in D3R revealed that bitopic PF ligands generally exhibited higher potency than the SM series ligands, with a susbtantial potency difference seen for the pair PF*_S,S_*-D*_S,R_* vs. SM-D*_S,R_*. This potency pattern is in line with scaffold specificity as PF and SM have higher affinity towards D3R and D2R respectively (Supplementary Table 3, [9, 10]). Further analysis of the pair PF*_S,S_*-D*_S,R_ vs.* SM-D*_S,R_* revealed higher potency of PF*_S,S_*-D*_S,R_* up to 35-fold in D2R, and up to 124-fold in D3R compared to SM-D*_S,R_*, across different transducers (except D3R-βarr1), supporting the role of the primary pharmacophore in linker-specific modulations. In particular, PF*_S,S_*-D*_S,R_*, showed the highest potency among PF series ligands in both D2R and D3R, across all transducers. On the other hand, SM-D*_S,R_*, exhibited the lowest potency within the SM series across all transducers. It is noteworthy that this comparison of PF*_S,S_*-D*_S,R_* vs. SM-D*_S,R_* revealed drastic potency differences, which are based solely upon the primary pharmacophore.The significant disparity in potency for these linker D ligands in D2R and D3R highlights the unique characteristics of D*_S,R_* linker as compared to other linkers [8]. It is also noteworthy that its diasteromer D*_R,S_* linker is favored in D2R-transducer activation.

With respect to E_max_ differences in D2R (Table 1), PF*_S,S_* and SM showed similar efficacy; whereas in D3R (Table 2), PF*_S,S_* exhibited considerably lower efficacy across several transducers (except βarr2), up to 96% less than SM. No significant difference in efficacy was apparent between PF*_S,S_*-A and SM-A in D2R, except for D2R-GoA, where there was a 50% increase in PF*_S,S_*-A E_max_ over SM-A. In D3R, PF*_S,S_*-A had ∼35% higher E_max_ compared to SM-A in GoA and Gi3, with an even greater 65% increase in βarr2. When evaluating other bitopic pairs, we observed at least 30% higher E_max_ for SM-based ligands in linker D pairs (i.e., SM-D*_S,R_*,over PF*_S,S_*-D*_S,R_* and SM-D*_R,S_* over PF*_S,S_*-D*_R,S_*) in D2R-βarr1 and D2R-βarr2. A tendency for higher efficacy of SM-C*_cis_* compared to PF*_S,S_*-C*_cis_* was noted for most transducers, with 115% higher efficacy in D2R-βarr1. No difference in efficacy greater than 30% was seen for other ligand pairs across all transducers. With regard to efficacy differences in D3R, nearly half of all ligands with identical linker pairs across all transducers showed a substantial E_max_ disparity (>30%), with PF-based ligands having higher efficacy than SM-based ligands. Among these, it is noteworthy that the pair PF*_S,S_*-D*_S,R_ vs.* SM-D*_S,R_* exhibited the most consistent E_max_ separation across all transducers, with differences ranging between 27-124%; in D3R-GoA and -GoB, both linker D-based pairs (i.e., PF*_S,S_*-D*_S,R_ vs.* SM-D*_S,R_*, PF*_S,S_*-D*_R,S_ vs.* SM-D*_R,S_*) showed E_max_ differences which ranged from 43-85%. In D3R-βarr2, a remarkable E_max_ difference of 93% was seen for the comparison of PF*_S,S_*-C*_trans_ vs.* SM-C*_trans_*. Additionally, in D3R-Gi1 and -Gi3, both linker B-based pairs (i.e., PF*_S,S_*-B*_cis_ vs.* SM-B*_cis_*, PF*_S,S_*-B*_trans_ vs*. SM-B*_trans_*) showed E_max_ differences ranging from 51-88%. Evaluating the transducer profile, in D3R-Gi1 and D3R-Gi3, all the bitopic PF ligands (Figure 3, L and N) led to a higher Emax response as compared to the corresponding pairs of SM ligands (Supplementary Figure S3, D and F).

Interestingly, we observed for linkers B, C, and D, consistent E_max_ within the same transducer regardless of linker chirality or primary pharmacophore (i.e., among four different ligands sharing the same linker type). For instance, when comparing ligands within Linker B (i.e., PF*_S,S_*- B*_cis_*, PF*_S,S_*-B*_trans_*, SM-B*_cis_*, SM-B*_trans_*), we observed minimal variation in E_max_ values within a specific transducer despite the significant differences apparent across various transducers. Specifically, in D2R, the average E_max_ values for these four ligands were as follows: 93-97% (GoA); 95-107% (GoB); 69-81% (Gz); 46-56% (Gi1); 70-86% (Gi3); 24-30% (βarr1); 96-106% (βarr2) (Table 1). This minimal E_max_ variation (i.e., < 20% variation) across ligands with the same linker type was also observed within Linker B in D3R across various transducers, including D3R-GoA, -βarr1, and -βarr2. Linker C ligands (i.e., PF*_S,S_*-C*_cis_*, PF*_S,S_*-C*_trans_*, SM-C*_cis_*, SM-C*_trans_*) also showed < 20% E_max_ variation in D2R and D3R-GoB and -Gz. Linker D ligands (i.e., PF*_S,S_*- D*_S,R_*, PF*_S,S_*-D*_R,S_*, SM-D*_S,R_*, SM-D*_R,S_*) showed < 20% E_max_ variation for many transducers in D2R. However, for most transducers in D3R, significant discrepancies in efficacy which primarily stem from modifications to the primary pharmacophore (i.e., E_max_ PF*_S,S_*-D*_S,R_* and PF*_S,S_*-D*_R,S_* > E_max_ SM-D*_S,R_* and SM-D*_R,S_*) were seen.

Overall, SM and bitopic SM ligands demonstrated slightly higher potency as well as higher efficacy (in partiular for β-arrestin recruitment) in D2R when compared to PF*_S,S_* and bitopic PF ligands within the same linker (except PF*_S,S_*-D*_S,R_*). Contrary to these results, in D3R, bitopic PF ligands surpassed bitopic SM ligands which share the same linker for both potency and efficacy across multiple transducers. These results emphasize the robust influence of the PF and SM primary pharmacophores on receptor selectivity within bitopic agonists. Nevertheless, linker type had an important role in agonist efficacy, particularly linkers B and C, impacting the activation of various transducers. Finally, the D*_S,R_* linker exerts a favorable change to PF*_S,S_* and an unfavorable potency change to SM in D3R- and D2R-transducer activation respectively.

### 5. Impact of chirality within the diastereomeric pair of bitopic ligands

In section 1, we examined the role of primary pharmacophore chirality as a determinant of potency and divergent behavior of diastereomers, including bitopic diastereomers with linker A. In the remaining bitopic ligands, linkers B and C have their relative stereochemistry resolved (i.e., *cis* and *trans* pairs for B; only one *cis*- and one *trans*- for C due to linker simmetry) while linker D has both its *trans*- stereosiomers resolved and their absolute configurations assigned (i.e., -D*_S,R_* and -D*_R,S_*). In this section, we analyzed the impact of chirality within each linker type by performing pairwise comparisons between bitopic diastereomers (e.g., PF*_S,S_*-B*_cis_* vs. PF*_S,S_*- B*_trans_*) to evaluate the pharmacological profile of each igand.

In comparisons of potency among diastereomer pairs, a pronounced separation was observed for linker D pairs (i.e., PF*_S,S_*-D*_S,R_*, vs. PF*_S,S_*-D*_R,S_* and SM-D*_S,R_* vs. SM-D*_R,S_*). PF*_S,S_*-D*_S,R_* showed 14-57-fold higher potency in D2R (Supplementary Figure S3, I-P) and 6-45-fold higher potency across different transducers in D3R (Figure 3, I-P) compared to PF*_S,S_*-D*_R,S_*. Interestingly, the equivalent pair of diastereomers with the SM scaffold (i.e., SM-D*_S,R_* vs. SM-D*_R,S_*) exhibited an inverse pattern in D2R (Supplementary Figure S3, A-H), with SM-D*_R,S_* being consistently more potent than SM-D*_S,R_* across different transducers.

Upon examining the effects of chirality on transducer profiles, we observed a pronounced impact on β-arrestin recruitment in both D2R and D3R. Specifically, across linker B and C diastereomers (i.e., PF*_S,S_*-B*_cis_* vs. PF*_S,S_*-B*_trans_*, SM-B*_cis_* vs. SM-B*_trans_*, PF*_S,S_*-C*_cis_* vs. PF*_S,S_*-C*_trans_*, and SM-C*_cis_* vs. SM-C*_trans_*), a minor, but consistent, potency separation was apparent in D2R- βarr1 (5-7-fold higher for *trans* than *cis*). In D3R-βarr1, similar consistency was not observed. However, notable differences in potency between diastereomers were evident, particularly in the comparison of SM-C*_cis_* vs. SM-C*_trans,_*, where the potency of *trans* was 1023-fold higher than that of *cis*. A similar disparity was seen for SM-B*_cis_* vs. SM-B*_trans_*, where the potency of *cis* was 525-foldhigher than that of *trans*.

Assessing efficacy, the vast majority of ligands exhibited differences of no more than 20% between diastereomeric pairs in both receptors. However, there are some notable differences: SM-D*_R,S_* showed consistently higher efficacy compared to SM-D*_S,R_* across all transducers in D2R and D3R, with E_max_ difference ranging between 4-47% in D2R (Table 1) and 16-37% in D3R (Table 2). In SM linker C diastereomers, SM-C*_cis_* showed higher E_max_ compared to SM-C*_trans_* by 116%, 290%, and 73% in D2R-βarr1, D3R-βarr1, and D3R-βarr2 respectively.

In short, chirality differences within PF diastereomers of linker D (i.e., PF*_S,S_*-D*_S,R_*, vs. PF*_S,S_*-D*_R,S_*) were found to significantly impact potency across all transducers in both D2R and D3R. While no significant differences in potency were observed for other diastereomeric pairs, this analysis revealed the influence of linker chirality on βarr1 recruitment. Interestingly, diastereomeric differences did not affect E_max_ responses for most pairs. Moreover, the impact of linker chirality on E_max_ appears to be receptor- and transducer-specific, with more prominent responses observed in D3R-Gi1, -Gi3, -βarr1, and -βarr2.

### 6. Kinetic analysis of potency and efficacy for SM and PF ligand series

The current study also aims to unravel the temporal aspect of functional outcomes in ligand potency and efficacy. By employing rigorous kinetic analyses, we analyzed ligands having stable or divergent kinetic readouts in an attempt to discover underlying trends. The activity of these ligands was assessed over a duration of 44 min; their kinetic characteristics were then analyzed by comparing the 10-min and 40-min time points. These two time points were specifically selected because the response typically stabilizes after an initial 10-min ligand incubation, and because a 30-min interval (i.e., 10 to 40 min) can be considered as a relevant time scale for the occurrence of molecular and cellular events.

When assessing potency over time, the reference agonist DA exhibited pEC_50_ values that were predominantly constant over time in both D2R (Figure 4, I-P) and D3R (Figure 5, G-L). DA potency variation between 10 min and 40 min was consitstently below 6-fold, with the exception of D2R-Gi2 (increase over 10-fold). In D2R-Gi2, an increase in potency of at least 30-fold was also observed for other ligands, including SM, SM-A, SM-D*_S,R_*, SM-D*_R,S_*, and PF*_S,S_*-D*_R,S_*. Furthermore, over time, there was a remarkable 52-fold potency decrease in PF*_S,S_* activation of D2R-Gi2. Despite this notable case, ligand potency remains stable for the majority of receptor-transducers activations over time.

**Figure 4.**
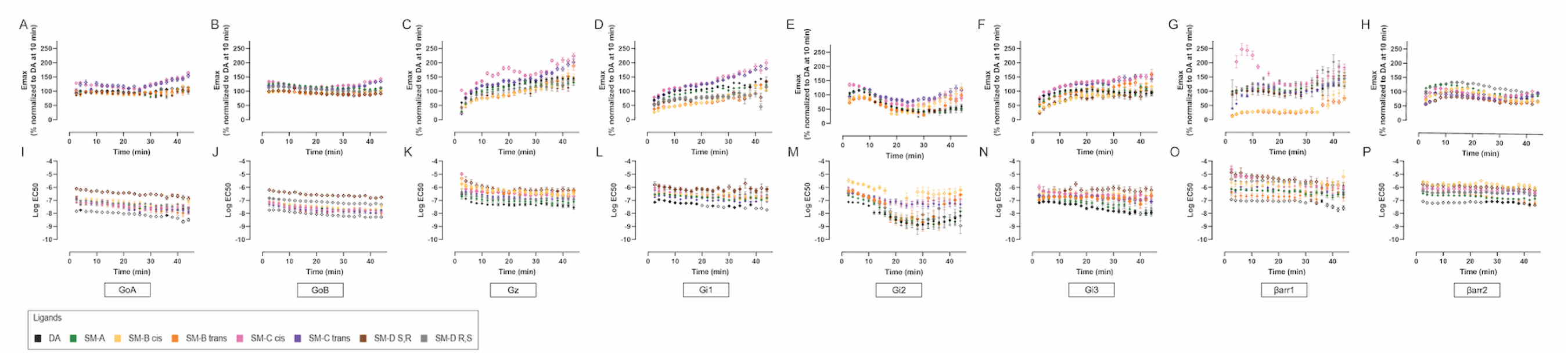
Kinetic profile of D2R-transducer activation for bitopic sumanirole ligands. Time-dependent plots of efficacy (A-H) and potency (I-P) changes for G protein activation (A-F, I-N) and β-arrestin recruitment (G-H, O-P) with measurements taken every two minutes, from 2 to 46 minutes post-ligand application. Data is presented as means ± SEM, normalized to DA at 10 min within bitopic SM series (n ≥ 4). Open symbols for data points denote *p < 0.05* compared to SM-A.

**Figure 5.**
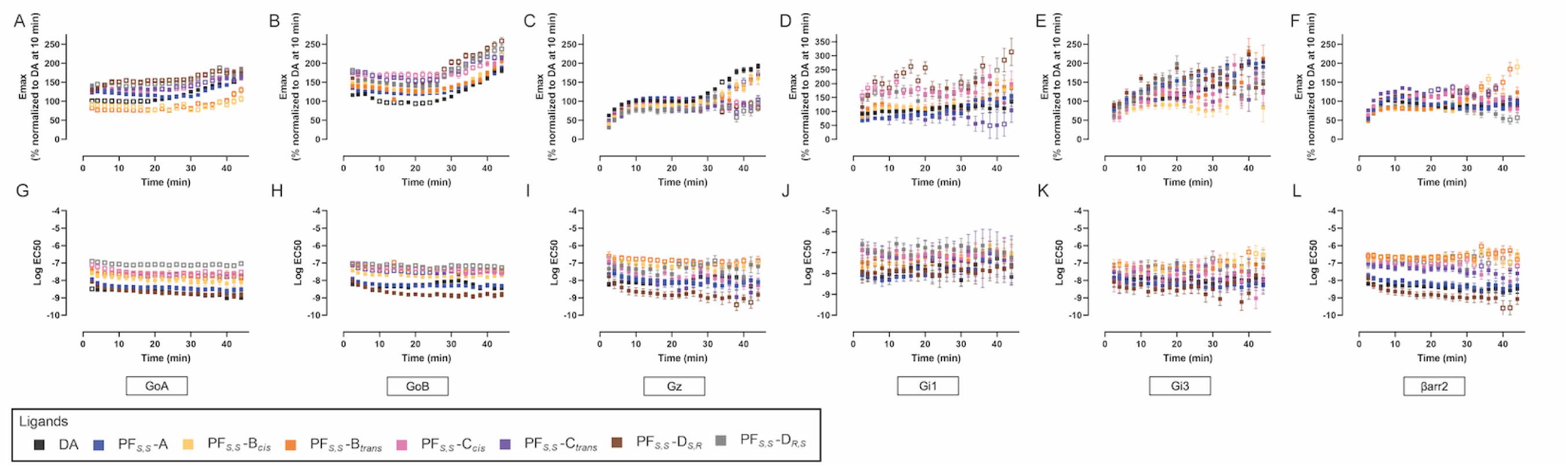
Kinetic profile of D3R-transducer activation for bitopic PF592,379 ligands. Time-dependent plots of efficacy (A-F) and potency (G-L) changes for G protein activation (A-E, G-K) and β-arrestin recruitment (F, I) with measurements taken every two minutes, from 2 to 46 minutes post-ligand application. Data is presented as means ± SEM, normalized to DA at 10 min within bitopic PF series (n ≥ 4). Open symbols for data points denote *p < 0.05* compared to PF-A.

When evaluating efficacy over time, in activation of G proteins, DA showed an E_max_ increase ranging between 8-43% in D2R (Figure 4, A-H) and 22-100% in D3R (Figure 5, A-F). Primary pharmacophore ligands (i.e., SM, PF*_S,S_* and PF*_R,S_*) as well as bitopic ligands strongly increased E_max_ over time for multiple G-proteins in D2R and D3R (except D2R-Gi2 which drastically decreased, up to 75% for most ligands). Across all ligands, D2R-Gi2 exhibits a distinctive kinetic profile which is characterized by a reduction in E_max_ over time. The kinetic profiles of primary pharmacophore ligands reveal a greater E_max_ change (either increase or decrease) over time in D3R compared to D2R that is consistent across all G-proteins. In particular, PF*_S,S_* and PF*_R,S_* showed a significant change in E_max_ over time (>30% increase) for Gz in D2R (Table 1 vs. Supplementary Table 1) and D3R (Table 2 vs. Supplementary Table 2), these same primary pharmacophores exhibited the most substantial increase in E_max_ across all D3R-G proteins (66-151%) over time, surpassing that of PF*_S,S_*-A (particularly in GoA) as seen in Kinetic plots (Supplementary Figure S7, A-F). Similarly, SM led to a change in E_max_ over time, primarily in D3R-G proteins (18-73%) (Supplementary Figure S8, A-F) and D2R-Gz (>30%) (Supplementary Figure S9, A-H). When analyzing the kinetic profile of bitopic ligands in D2R, all bitopic ligands regardless of scaffold or linker exhibited increased E_max_ over time for D2R-Gz and -Gi1. This was significant (>30%) for the majority of ligands, of which PF*_S,S_*-B*_cis_*, SM-B*_cis_*, and SM-C*_cis_* exhibited still greater increase (>70%). Furthermore, an increase in E_max_ over time greater than 30% was observed in D2R-GoA when activated by bitopic PF ligands which featured linkers C and D (i.e., PF*_S,S_*-C*_cis_*, PF*_S,S_*-C*_tran_*, PF*_S,S_*-D*_S,R_*, PF*_S,S_*-D*_R,S_*); this was also observed in D2R-Gi3 upon activation by bitopic PF ligands with linkers B and D (i.e., PF*_S,S_*-B*_cis_*, PF*_S,S_*-B*_trans_*, PF*_S,S_*-D*_S,R_*, PF*_S,S_*-D*_R,S_*) (Supplementary Figure S5, A-H), and bitopic SM ligands with linkers B and C (SM-B*_cis_*, SM-B*_trans_*, SM-C*_trans_*, SM-C*_cis_*) (Figure 4, A-H). In D3R, >30% increase in E_max_ was observed over time in D3R-Gz and -Gi1 upon activation by all ligands with linkers A and B (25-95%); substantial increase in E_max_ was also apparent in D3R-GoB upon activation by all bitopic PF ligands, of which those with linker D (i.e. PF*_S,S_*-D*_S,R_*, PF*_S,S_*-D*_R,S_*) exihibited 76-91% range, as well as in D3-Gi3 where a 78-121% increase was observed when activated by the specific ligands PF*_S,S_*-A, PF*_S,S_*-B*_trans_*, SM-B*_cis_*, SM-C*_trans_*, and SM-C*_cis_* (Figure 5, A-F ands Supplementary Figure S4, A-F).

With regard to the kinetic profile of β-arrestin recruitment in D2R-βarr1, a significant increase in E_max_ over time was observed for various ligands, including SM (102%), SM and PF ligands with linker B (35-208%) and SM ligands with linker D (45-78%) (Table 1 vs. Supplementary Table 1 and Table 2 vs. Supplementary Table 2). Conversely, in D2R- βarr2, there was a reduction in E_max_ up to 42% for all ligands tested. On the other hand, for D3R, the βarr1 and βarr2 responses follow identical trends of E_max_ change (either increase or decrease over time) for most bitopic ligands, including PF*_S,S_*-A, SM-B*_cis_*, SM-B*_trans_* and SM-C*_cis_* with 13-145% decrease and PF*_S,S_*-B*_cis_*, PF*_S,S_*-B*_trans_*, and SM-C*_trans_*, with 38-198% increase from 10 to 40 min.

The clear divergence in kinetic profiles of D2R compared to D3R, when activated by bitopic ligands, is observed in GoB activation by PF*_S,S_*-D*_R,S_* and SM-D*_S,R_*, for which the increase in E_max_ over time is 62-91% in D3R, but not significant in D2R, and in Gi1 activation by SM-C*_cis_* and PF*_S,S_*-D*_S,R_*, which conversely show an increase in E_max_ over time of 64-75% in D2R but no significant change in D3R. Interestingly, in βarr2 recruitment by PF*_S,S_*-B*_cis_* and PF*_S,S_*-B*_trans_*, we observed a 31-34% decrease in E_max_ for D2R, but a trend in the opposite direction for D3R, where a 38-67% increase in E_max_ was observed. Notably, E_max_ differences between diastereomeric pairs tend to increase over time, particularly for PF ligands with linker B (i.e., PF*_S,S_*-B*_cis_* vs. PF*_S,S_*-B*_trans_*) in D2R and D3R, and for most diasteomeric pairs in D3R-Gi1 and D3R-Gi3.

In summary, the majority of ligands maintained stable potency over time across most receptor-transducer combinations. However, there was significant variability in ligand efficacy over time. Increasing E_max_ over time was particularly pronounced for Gz, Gi1, and Gi3 in both D2R and D3R, across various ligands, while declining E_max_ was exclusive to D2R-βarr2, D2R-Gi2, and D3R-Gi2 across all ligands. Regardless of SM or PF series, most ligands demonstrated a trend of increased efficacy over time across most transducers (except Gi2 and βarr2 in D2R and D3R), a pharmacological behavior unlikely to be associated with the primary pharmacophore.

### 7. Comparison of bias factors at 10 and 40 min for D2R and D3R

Bias factor calculation allows assessment of preferential activation and/or recruitment among 8 transducers by an agonist in D2R and D3R separately. The bias factor, calculated as described in the methods section, uses DA as a reference agonist and GoA as a reference transducer, represented as ΔΔ for a specific timepoint. If ΔΔ is negative, it indicates a preference for GoA activation, referred to as ’bias towards GoA.’ Conversely, positive ΔΔ means preferential activation of the compared transducer over GoA, implying ’bias away from GoA.’ We set a significance threshold for ΔΔ values −1 and 1 (i.e., ΔΔ values <-1 and >1 are considered significantly biased.) Multicomparison data analyses at the 10 min and 40 min time points were conducted to discern unique agonist profiles. Additionally, a temporal comparison of bias factors was derived by calculating the difference between the bias factors at 10 and 40 min.

When assessing bias in D2R, within the SM series, no significant bias towards any particular D2R-transducer among SM and bitopic SM ligands was seen at 10 min. However, a notable bias away from D2R-βarr1 was observed in the majority of bitopic SM ligands, with SM-B*_cis_*, SM-C*_cis_* and SM-D*_R,S_* having the highest ΔΔ values (ranging between −0.63 and −0.78) (Figure 6, A-D). At 40 min, both bitopic SM-B ligands exhibited significant bias away from D2R-βarr1, with ΔΔ −0.99 for SM-B*_cis_* and −1.09 for SM-B*_trans_*, a trend also observed for PF ligands with same linkers (i.e., PF*_S,S_*-B*_cis_* and PF*_S,S_*-B*_trans_*) (Figure 6, E-H). Additionally, SM-B ligands exhibited bias away from D2R-Gi2, with substantial ΔΔ of −1.36 for SM-B*_trans_*, as opposed to the SM-A and SM-D ligands which showed a tendency for bias towards D2R-Gi2 (with ΔΔ ranging between 0.69 and 0.82) at the later timepoint.

**Figure 6.**
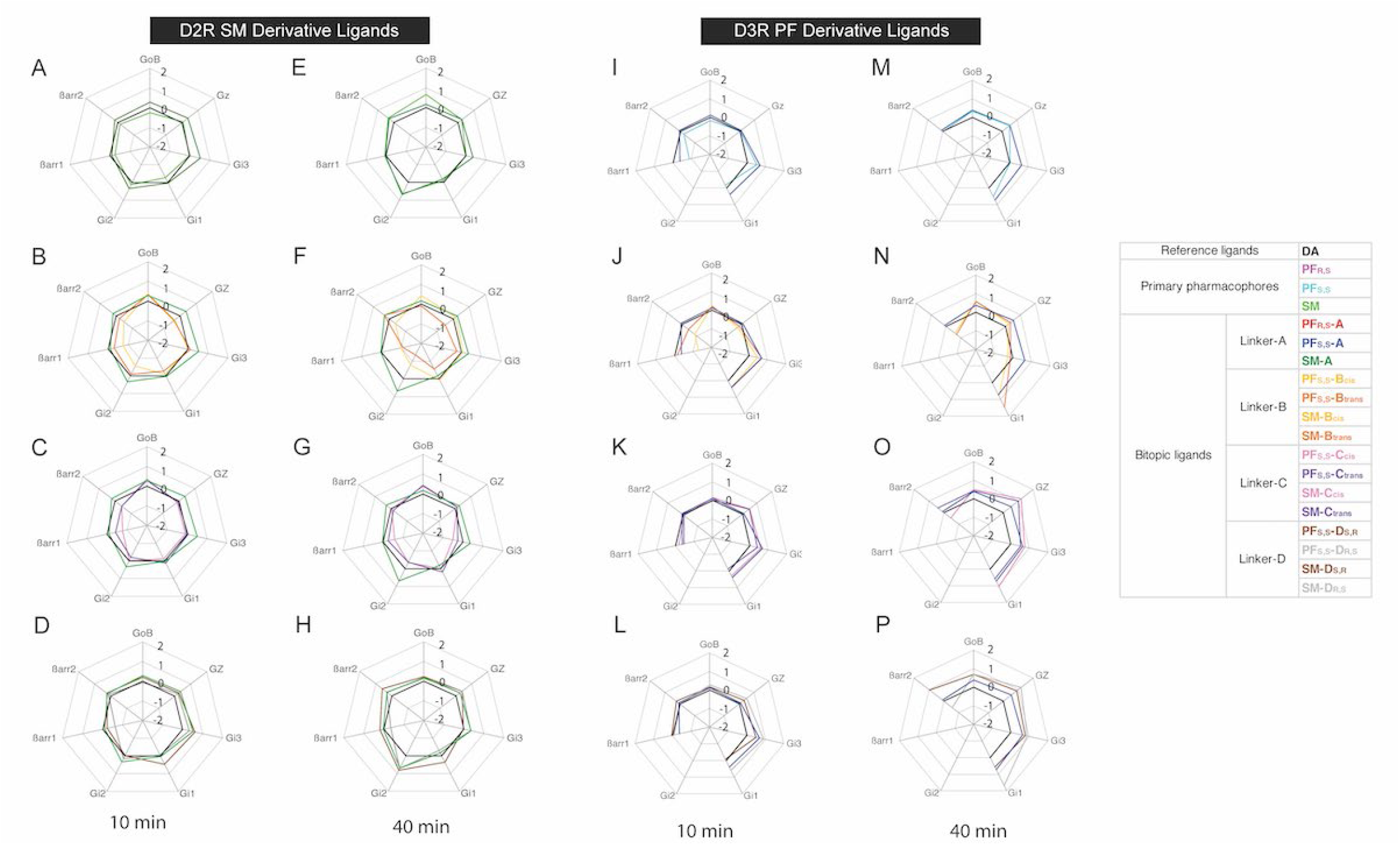
Bias factor profile of D2R- and D3R-transducer activation by bitopic sumanirole and PF592,379 ligands. Transducer activation potencies and efficacies in Tables 1 and 2 at 10 min and Supplementary Tables 1 and 2 at 40 min were used to calculate bias factors, plotted in spider charts on a logarithmic scale with DA as a reference agonist. D2R SM-derivative ligands (A-H), D3R PF592,379-derivative ligands (I-P) at 10 min (A-D and I-L) and 40 min (E-H and M-P).

Analysis of the PF series indicated no significant transducer bias at 10 min, except for PF*_S,S_*-B*_cis_* which showed a significant bias away from D2R-βarr1 (ΔΔ = −1.0) (Supplementary Figure 6, A-D). At 40 min, both PF-B ligands (i.e. PF*_S,S_*-B*_cis_* and PF*_S,S_*-B*_trans_*) showed a trend towards bias for D2R-βarr2 (ΔΔ = 0.65 and 0.61, respectively) in contrast with D2R-βarr1 (ΔΔ = −0.58 and −0.87, respectively) (Supplementary Figure 6, E-H), evidencing a considerable bias discrepancy between the two β-arrestin subtypes. Additionally, PF*_S,S_* exhibited a trend of bias towards D2R- βarr2 (ΔΔ = 0.65) and -Gz (ΔΔ = 0.55). Although the ΔΔ values were below 1, this trend was consistently observed across all bitopic PF ligands at 40 min. Specifically, PF ligands with linkers A and B showed bias towards D2R-βarr2, while PF ligands with linkers C and D showed bias towards D2R-Gz. Furthermore, there was remarkable bias towards D2R-Gi2 activation by PF*_S,S_*-D*_R,S_* (ΔΔ = 1.94), in stark contrast to its diastereomer PF*_S,S_*-D*_S,R_* (ΔΔ = −0.95).

When evaluating bias factors in D3R within the SM series, we observed considerable bias away from β-arrestin recruitment (i.e., towards GoA) at 10 min. Specifically, bias away from D3R- βarr1 was observed for most SM ligands, with ΔΔ values up to −1.26, except for SM-B*_cis_*, which showed opposite bias directionality (ΔΔ = 1.3). Bias away from D3R-βarr2 was also observed for SM and SM ligands with linkers B and C, with ΔΔ values as low as −1.53. Additionally, bias towards other G-proteins was present, for example bias towards D3R-Gi3 with SM-D*_R,S_* (ΔΔ = 1.06), and bias towards D3R-Gi1 with SM-B*_cis_* (ΔΔ = 1.28) relative to D3R-GoA at 10 min. The bias observed in D3R-Gi1 is indicative of a notable (>2) bias difference between SM-B*_cis_* (ΔΔ = 1.28) and its diastereomer, SM-B*_trans_* (ΔΔ = −1.16) (Supplementary Figure 6, I-L). At 40 min, bias towards D3R-Gi1 becomes more significant for all SM ligands, particularly for SM and SM ligands with linkers B and D, with ΔΔ values ranging between 1.09 and 1.95. Notably, bias differences between D3R-Gi1 vs. D3R-βarr2 exibited ΔΔ values above 2 for some bitopic ligands, including for SM-B*_cis_*, which exhibited a bias difference of 2.8 at 10 min, and 3.14 at 40 min (Supplementary Figure 6, M-P) for these transducers.

Analysis of the PF series in D3R at 10 min revealed that PF*_S,S_* and all bitopic PF ligands were consistently biased away from D3R-βarr1 recruitment, coupled with bias towards D3R-Gi3 activation at this time point. While PF*_S,S_*-B*_cis_* was the only ligand that showed a considerable bias towards D3R-βarr1 (ΔΔ = −1.26), the difference in bias between D3R-Gi3 and D3R-βarr1 was particularly significant for PF*_S,S_*, PF*_S,S_*-A and PF*_S,S_*-B*_cis_*, with differences of 1.38, 1.02 and 1.72, respectively (Figure 6, I-L). At 40 min, the D3R-Gi3 bias was sustained for most bitopic PF ligands except PF-B. Furthermore, other G proteins, including Gz and Gi1, were preferentially activated by PF ligands relative to GoA at later timepoints. This biased activation was significant in D3R-Gz for PF*_S,S_*-C*_cis_*, PF*_S,S_*-C*_trans_*, and PF*_S,S_*-D*_R,S_*, with ΔΔ values within the 1.01-1.2 range. Additionally, in D3R-Gi1, PF*_S,S_*-B*_trans_*, PF*_S,S_*-C*_cis_* and PF*_S,S_*-D*_R,S_* showed significant bias, with ΔΔ values ranging from 1.06 to 1.69. Notably, for D3R-Gi1, a clear bias separation between PF diastereomers was observed, particularly those with linkers B and D (Figure 6, M-P).

In summary, our findings reveal a consistent trend for bias away from βarr1 for most bitopic SM and PF agonists in both D2R and D3R. While bias for a certain signal transducer may not be apparent at early time points in D2R, the bias factor becomes significant over time. Additionally, for various ligands, a bias towards Gz and Gi2 was observed in D2R at 40min, while a bias towards Gz, Gi1 and Gi3 was observed in D3R at 10 and 40 min. Overall, our findings provide insight into transducer activation profiles across SM and PF bitopic ligand series. For instance, a bias away from βarr1 recruitment was observed in D2R but not D3R, particularly for linker B ligands of the PF and SM series, but not for ligands with other linker types. This underscores the potential role of the cyclopentyl linker (i.e., linker B) in lowering βarr1 recruitment to D2R, irrespective of the primary pharmacophore. Gi1 activation bias was observed in D3R but not in D2R for both SM and PF ligands, regardless of the linker. More pronounced Gi1 activation bias was observed in D2R for SM and SM ligands with linkers B and D, with bias factors that increased over time. Gz activation bias was observed in both D2R and D3R for most ligands at 10 min, and significantly increased at 40 min. However, this bias was was absent in linker B PF ligands, an observation suggesting relevance of the cyclopentyl linker in bitopic PF ligands for suppression of Gz activation.

The current study provides a unique, comprehensive analysis of transducer activation profile and SAR for linker type in D2R- and D3R-selective bitopic agonists. We found that all linkers A-D showed differences in characteristics, whether sutble or significant. Notably, we observed a potency shift of Linker A for PF*_S,S_*-A vs. PF*_R,S_*-A, kinetic bias in Linker B for SM-B and PF-B, and enhanced potency in Linker C for SM-C over SM-A. Linker D showed enhanced potency of PF*_S,S_*-D over PF*_S,S_*-A, but had a significant difference within the diastereomers (i.e., PF*_S,S_*-D*_S,R_*, vs. PF*_S,S_*-D*_R,S_*). Ligand linker structure plays a role in not only favorable orientation of the secondary pharmacophore within the binding pocket, but also in formation of interactions within the vestibule between the orthosteric and allosteric sites; thus, is a vital component of receptor activation profiles. Our analysis revealed that different linker structures, including chirality in diastereomers, directly shape transducer activation profiles and their changes over time. Therefore, alternative linkers within bitopic ligands should be considered in drug design for desired receptor activation profiles, aside from their implications in metabolic stability. Beyond the improved activation profiles of compounds which are provided by this study, our extensive SAR work can serve as a template for future medicinal chemistry and molecular pharmacology efforts within the current landscape of in-depth transducer characterization and biased agonism.

## Author Contributions

HY and AS designed the experiments. AS, RG, JF, and WZ carried out the experiments, and performed the data analysis with HY and YJJ. KHL and LS performed the docking study. AB, FOB, and AHN performed the chemical synthesis. AS, RG, and HY prepared the initial manuscript, with contributions from all the authors in finalizing the manuscript.

## Acknowledgements

Translational Analytical Core at NIDA-IRP for HRMS-MS/MS analysis of the tested compounds. The work was supported by Northeastern University Startup Funds and Tier1 Intramural Grant (HY) and by the National Institute on Drug Abuse intramural research program (LS, AHN).

---

**Supplementary Figure 1.**
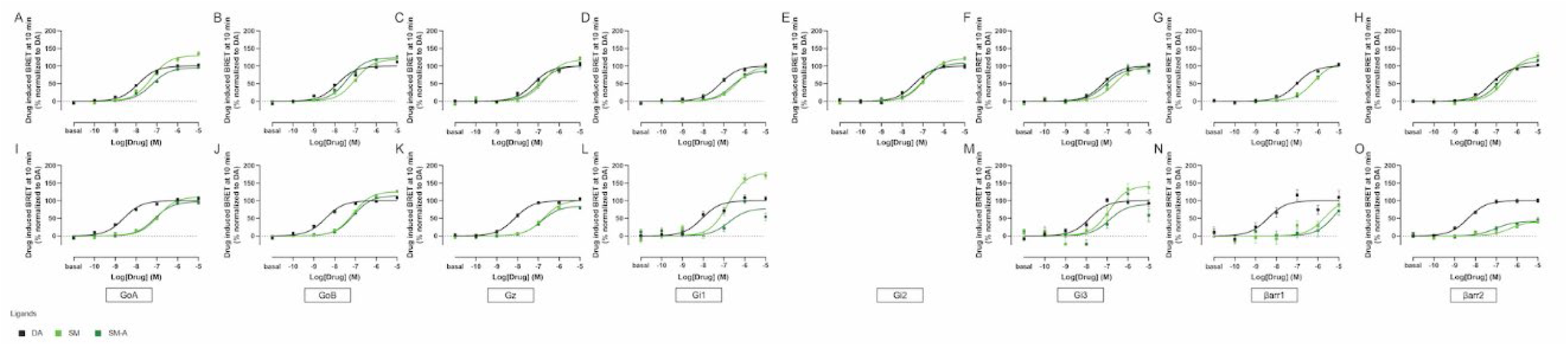
D2R- and D3R-transducer activation profile of bitopic sumanirole ligands with butyl linker at 10 minutes. Concentration-response curves for bitopic SM-ligands in D2R Gα-subtype activation (A-F), and β-arrestin-subtype recruitment (G-H). Concentration-response curves for bitopic PF-ligands in D3R Gα-subtype activation (I-M), and β-arrestin-subtype recruitment (N-O). Curves are presented as a percentage of the maximal response of DA with means ± SEM (n ≥ 4).

**Supplementary Figure 2.**
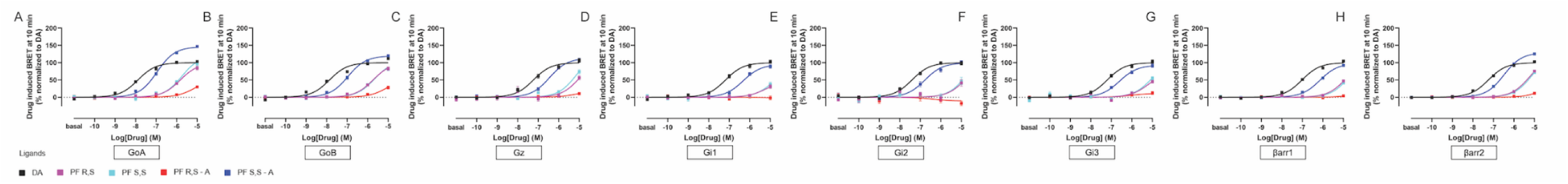
D2R-transducer activation profile of bitopic PF592,379 ligands with butyl linker at 10 minutes. Concentration-response curves of activation BRET between Gα-subtype-RLuc and Gγ-Venus (A-F), and D3R-RLuc and β-arrestin-subtype-Venus (G-H). Curves are presented as a percentage of the maximal response of DA with means ± SEM (n ≥ 4).

**Supplementary Figure 3.**
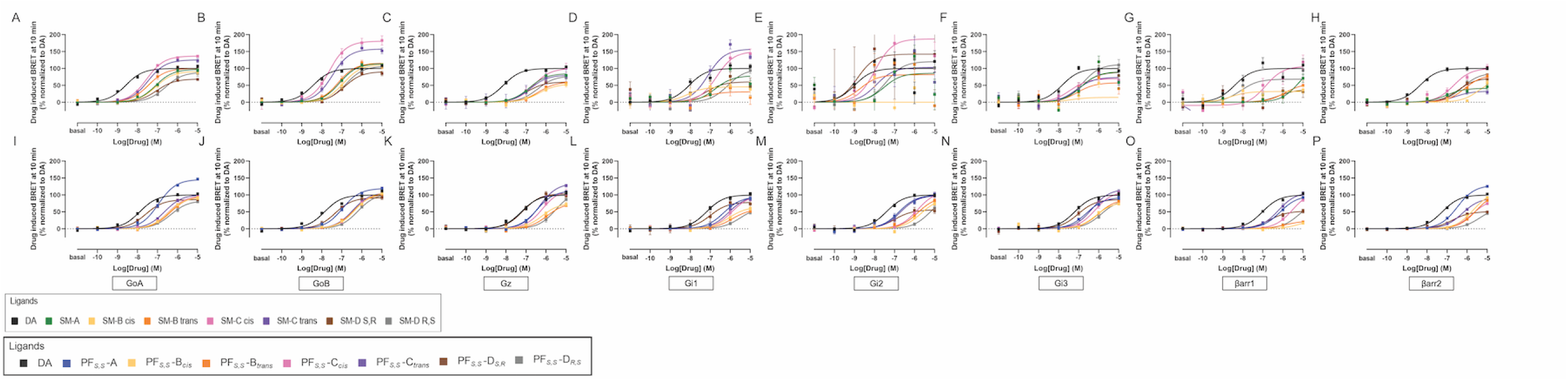
D2R- and D3R-transducer activation profile of bitopic PF592,379 and sumanirole ligands with various linkers at 10 minutes. Concentration-response curves for bitopic PF-ligands in D2R Gα-subtype activation (A-F), and β-arrestin-subtype recruitment (G-H). Concentration-response curves for bitopic SM-ligands in D3R Gα-subtype activation (I-N), and β-arrestin-subtype recruitment (O-P). Curves are presented as a percentage of the maximal response of DA with means ± SEM (n ≥ 4).

**Supplementary Figure 4.**
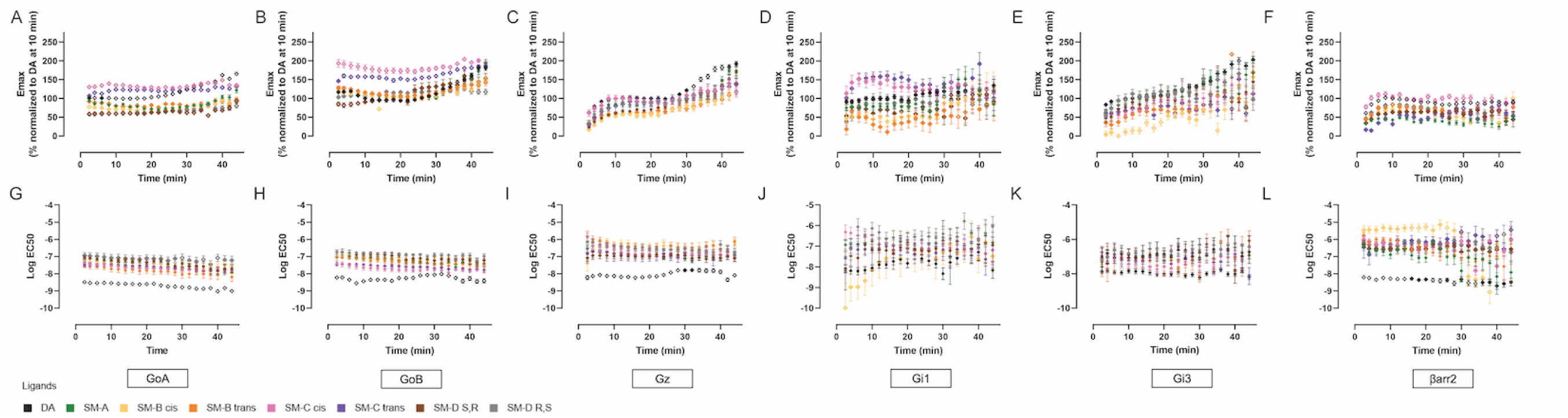
Kinetic profile of D3R-transducer activation for bitopic sumanirole ligands with various linkers. Time-dependent plots of efficacy (A-F) and potency (G-L) changes for G protein activation (A-E, G-K) and β-arrestin recruitment (F, I) with measurements taken every two minutes, from 2 to 46 minutes post-ligand application. Data is presented as means ± SEM, normalized to DA at 10 min within bitopic SM series (n ≥ 4). Open symbols for data points denote *p < 0.05* compared to SM-A.

**Supplementary Figure 5.**
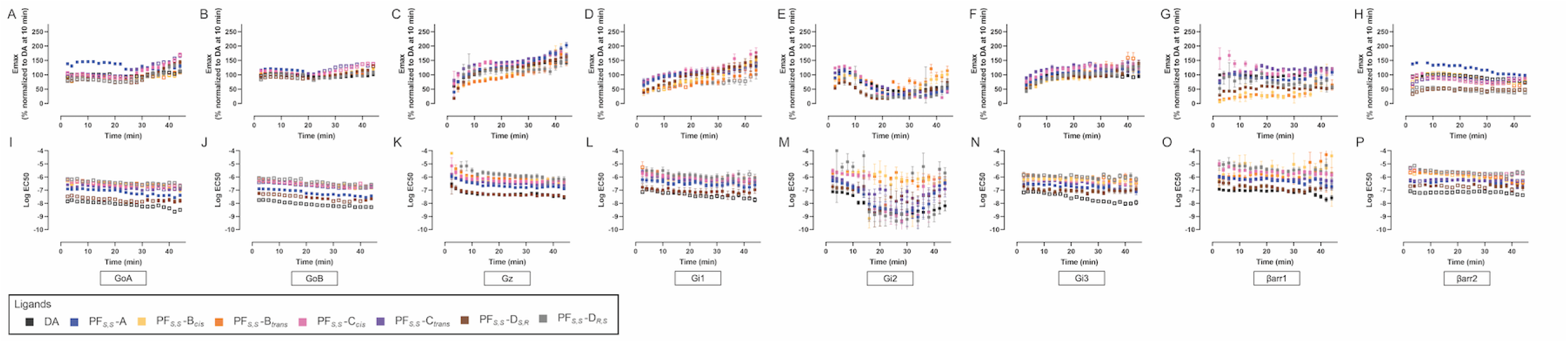
Kinetic profile of D2R-transducer activation for bitopic PF592,379 ligands with various linkers. Time-dependent plots of efficacy (A-H) and potency (I-P) changes for G protein activation (A-F, I-N) and β-arrestin recruitment (G-H, O-P) with measurements taken every two minutes, from 2 to 46 minutes post-ligand application. Data is presented as means ± SEM, normalized to DA at 10 min within bitopic PF series (n ≥ 4). Open symbols for data points denote *p < 0.05* compared to PF-A.

**Supplementary Figure 6.**
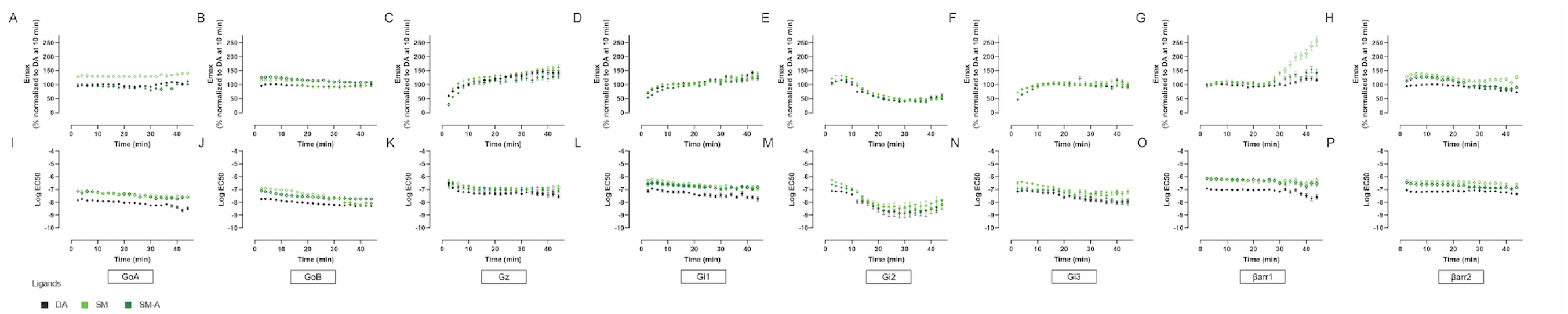
Kinetic profile of D2R-transducer activation for bitopic sumanirole ligands with butyl linker. Time-dependent plots of efficacy (A-H) and potency (I-P) changes for G protein activation (A-F, I-N) and β-arrestin recruitment (G-H, O-P) with measurements taken every two minutes, from 2 to 46 minutes post-ligand application. Data is presented as means ± SEM, normalized to DA at 10 min within bitopic SM series (n ≥ 4). Open symbols for data points denote *p < 0.05* compared to DA.

**Supplementary Figure 7.**
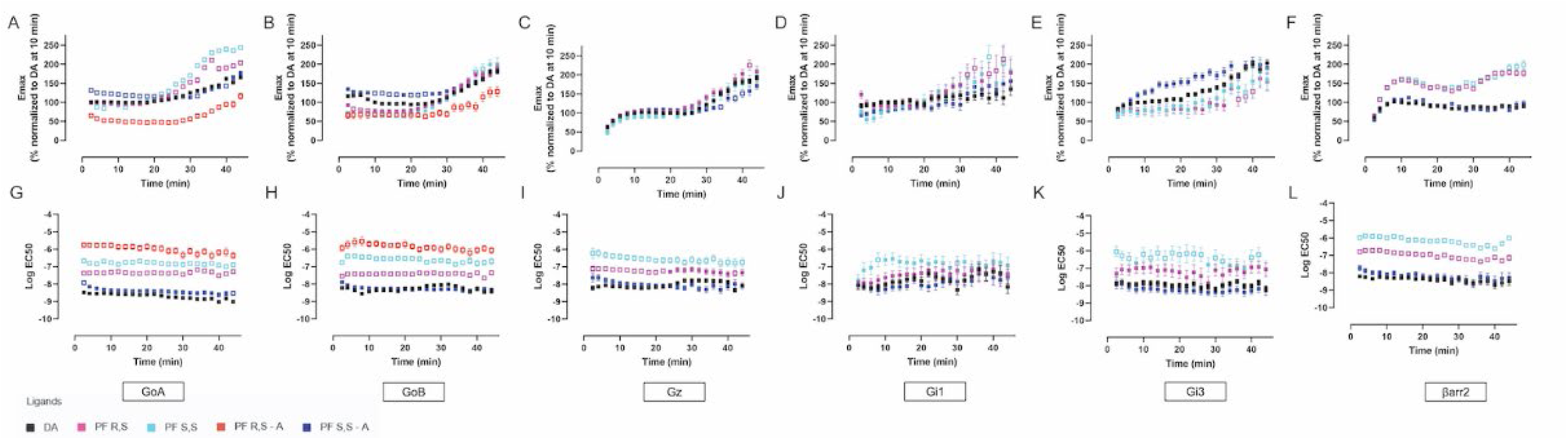
Kinetic profile of D3R-transducer activation for bitopic PF592,379 ligands with butyl linker. Time-dependent plots of efficacy (A-F) and potency (G-L) changes for G protein activation (A-E, G-K) and β-arrestin recruitment (F, I), with measurements taken every two minutes, ranging from 2 to 46 minutes post-ligand application. Data is presented as means ± SEM, normalized to DA at 10 min within bitopic PF series (n ≥ 4). Open symbols for data points denote *p < 0.05* compared to DA.

**Supplementary Figure 8.**
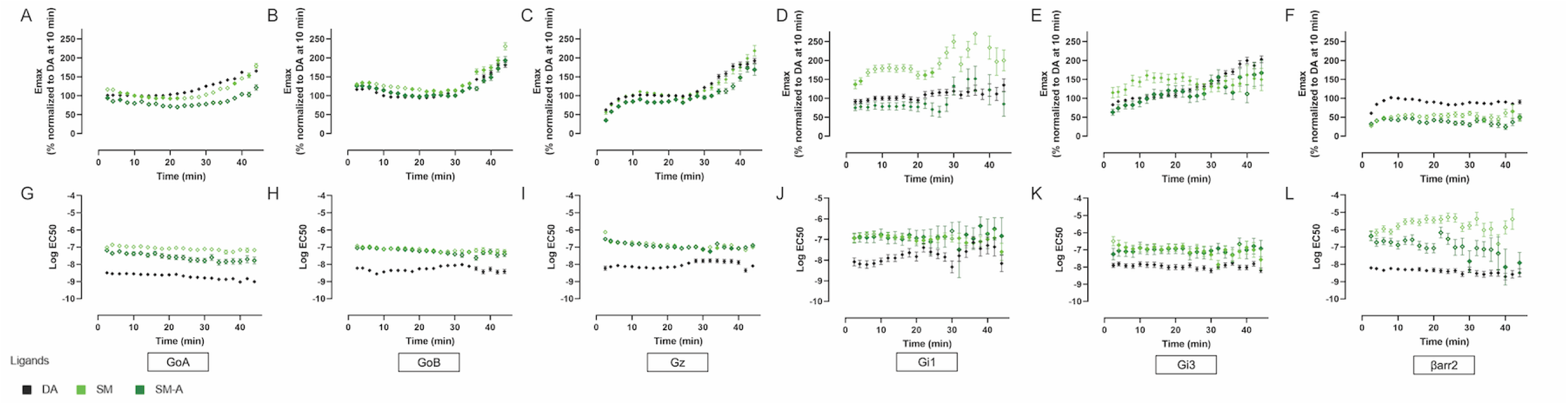
Kinetic profile of D3R-transducer activation for bitopic sumanirole ligands with butyl linker. Time-dependent plots of efficacy (A-F) and potency (G-L) changes for G protein activation (A-E, G-K) and β-arrestin recruitment (F, I) with measurements taken every two minutes, from 2 to 46 minutes post-ligand application. Data is presented as means ± SEM, normalized to DA at 10 min within bitopic SM series (n ≥ 4). Open symbols for data points denote *p < 0.05* compared to DA.

**Supplementary Figure 9.**
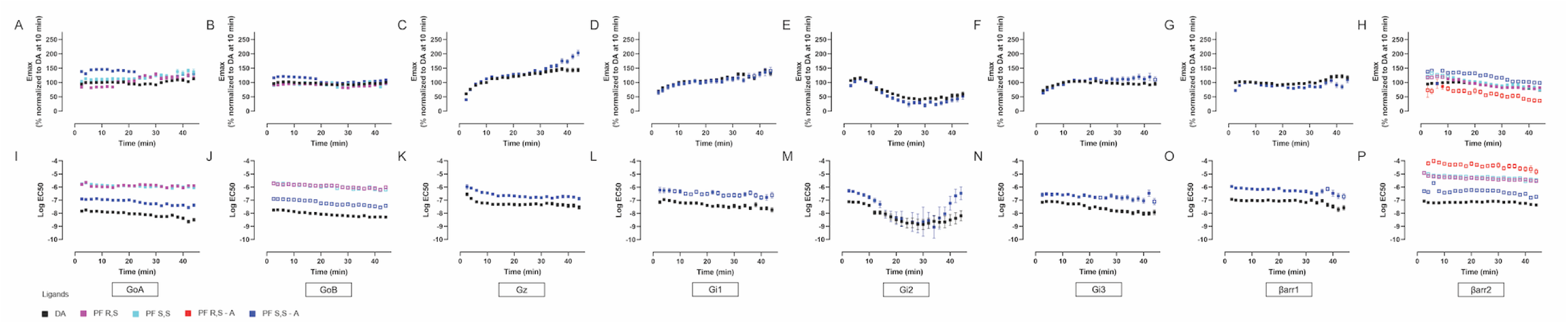
Kinetic profile of D2R-transducer activation for bitopic PF592,379 ligands with butyl linker. Time-dependent plots of efficacy (A-H) and potency (I-P) changes for G protein activation (A-F, I-N) and β-arrestin recruitment (G-H, O-P) with measurements taken every two minutes, from 2 to 46 minutes post-ligand application. Data is presented as means ± SEM, normalized to DA at 10 min within bitopic PF series (n ≥ 4). Open symbols for data points denote *p < 0.05* compared to DA.

**Supplementary Figure 10.**
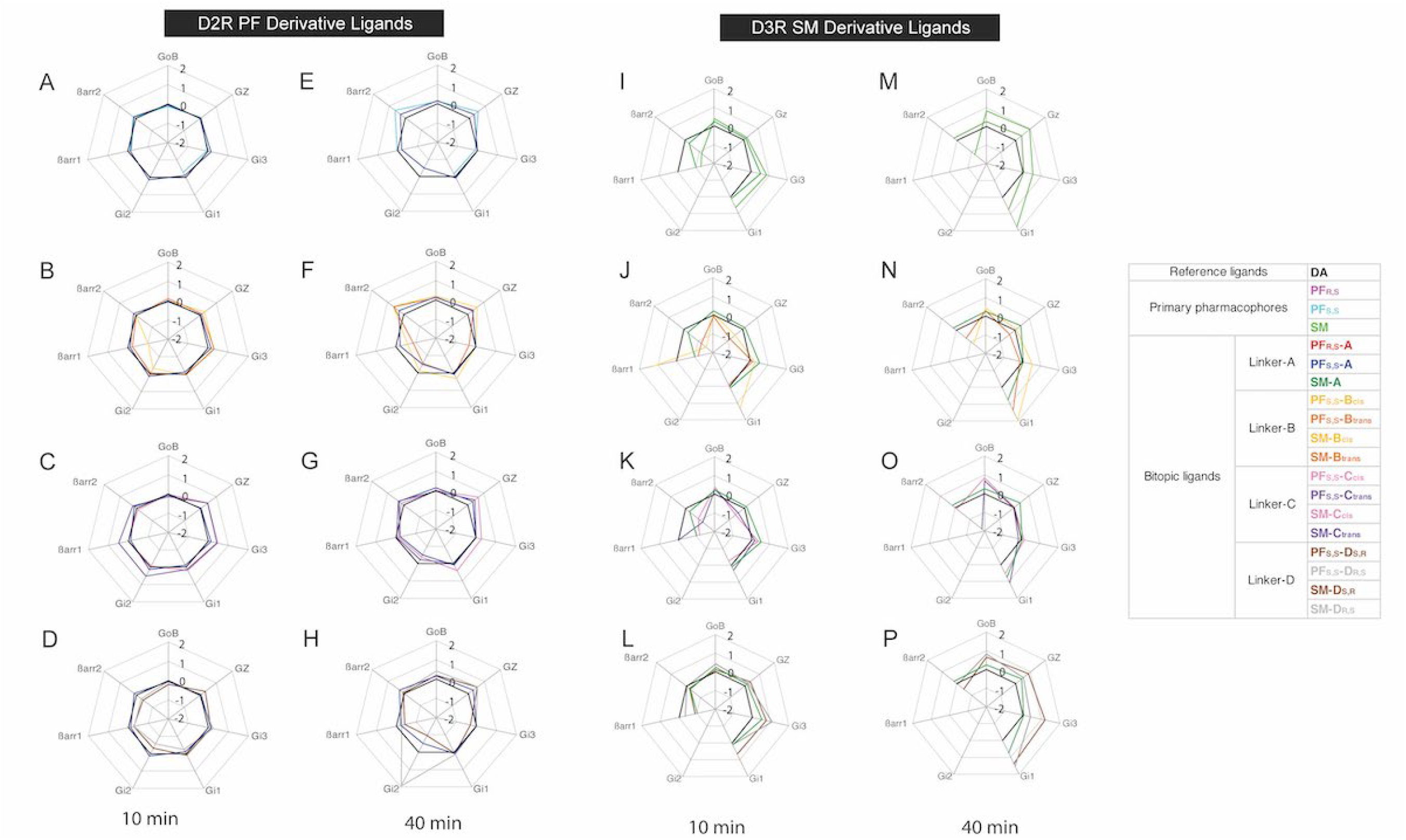
Bias factor profile of D2R- and D3R-transducer activation by bitopic sumanirole and PF592,379 ligands. Transducer activation potencies and efficacies in Tables 1 and 2 at 10 min and Supplementary Tables 1 and 2 at 40 min were used to calculate bias factors. They are plotted in spider charts on a logarithmic scale with DA as a reference agonist. D2R SM-derivative ligands (A-H), D3R PF592,379-derivative ligands (I-P) at 10 min (A-D and I-L) and 40 min (E-H and M-P).

**Supplementary Table 1.**
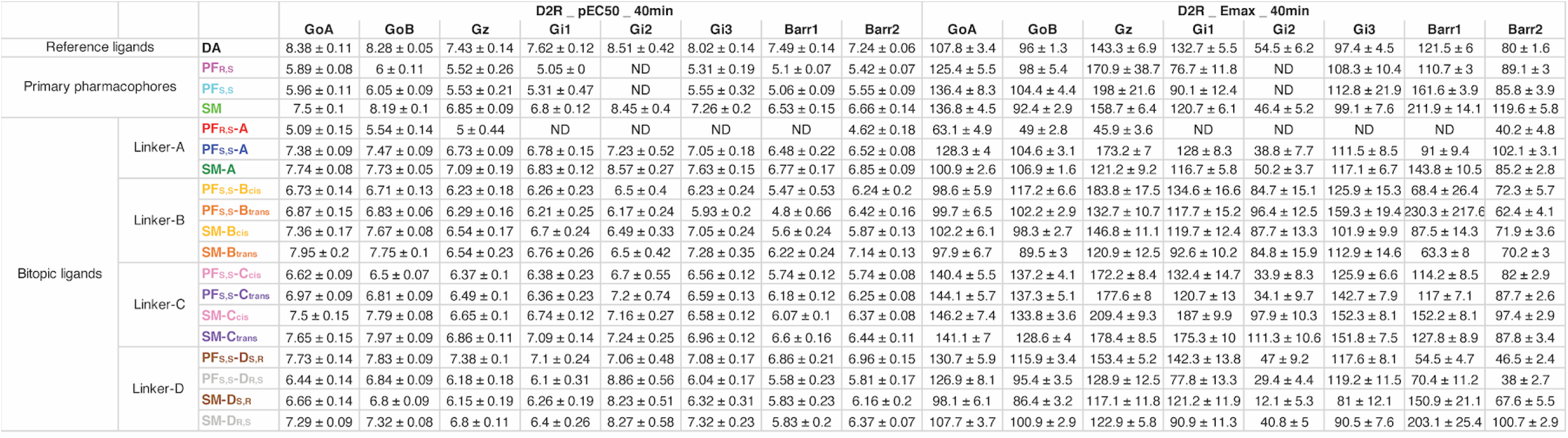
– Potency and efficacy of bitopic sumanirole and PF592,379 ligands for D2R activities at 40 min.

**Supplementary Table 2.**
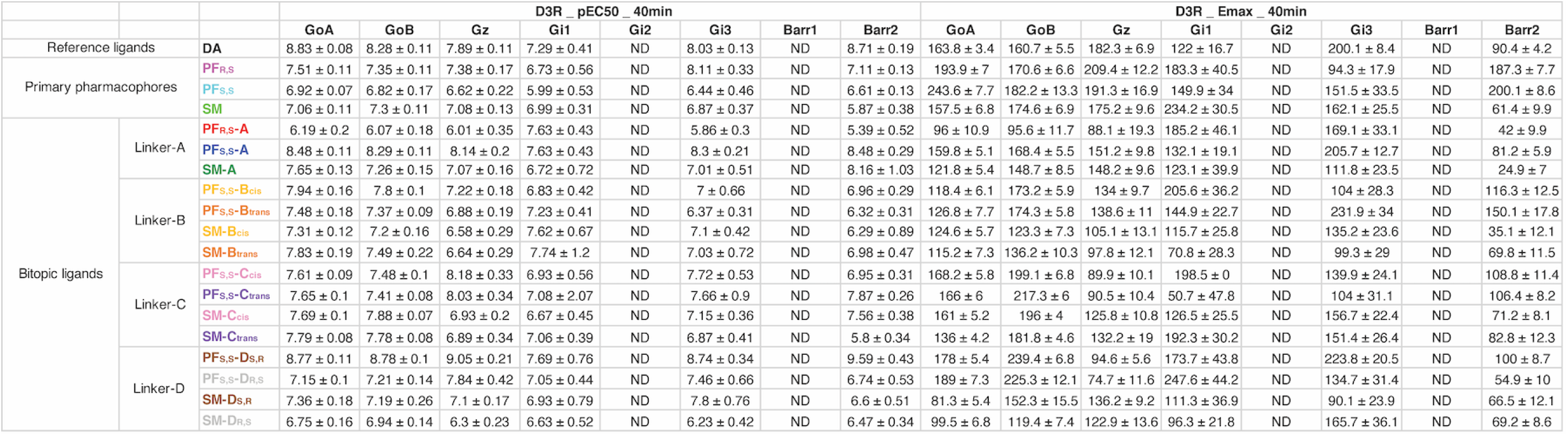
– Potency and efficacy of bitopic sumanirole and PF592,379 ligands for D3R activities at 40 min.

**Supplementary Table 3.** – Radioligand competition binding for D2R and D3R. [^3^H]-(*R*)-(+)-7-OH-DPAT radioligand binding assays performed on HEK293 cells stably expressing hD2LR and hD3R, following the previously reported protocols. Each *K_i_* value represents the arithmetic mean ± S.E.M of at least three independent experiments, each performed in triplicate. ^a,b,c^previously reported [7, 8, 10]

| Compound | D<sub>2</sub>R K<sub>i</sub> ± SEM (nM) | D<sub>3</sub>R K<sub>i</sub> ± SEM (nM) |
| --- | --- | --- |
| <sup>a</sup>SM | 80.6 ± 15.4 | 784 ± 131 |
| <sup>b</sup>PF<sub>R,S</sub> | 1510 ± 175 | 80.3 ± 13.3 |
| <sup>b</sup>PF<sub>S,S</sub> | 1320 ± 197 | 625 ± 118 |
| <sup>a</sup>SM-A | 15.7 ± 2.08 | 82.7 ± 13 |
| <sup>c</sup>PF<sub>R,S</sub>-A | 5220 ± 735 | 6470 ± 675 |
| <sup>c</sup>PF<sub>S,S</sub>-A | 134 ± 21.6 | 5.96 ± 0.477 |
| <sup>b</sup>SM-B<sub>cis</sub> | 2.03 ± 0.190 | 19.5 ± 0.450 |
| <sup>b</sup>SM-B<sub>trans</sub> | 4.58 ± 0.671 | 77.1 ± 10.6 |
| <sup>b</sup>PF<sub>S,S</sub>-B<sub>cis</sub> | 80.8 ± 11.0 | 5.72 ± 0.744 |
| <sup>b</sup>PF<sub>S,S</sub>-B<sub>trans</sub> | 115 ± 5.30 | 22.6 ± 2.01 |
| <sup>b</sup>SM-C<sub>cis</sub> | 5.73 ± 0.850 | 15.5 ± 2.48 |
| <sup>b</sup>SM-C<sub>trans</sub> | 11.9 ± 1.40 | 49.3 ± 4.30 |
| <sup>b</sup>PF<sub>S,S</sub>-C<sub>cis</sub> | 110 ± 9.51 | 31 ± 2.70 |
| <sup>b</sup>PF<sub>S,S</sub>-C<sub>trans</sub> | 86.2 ± 18.7 | 21 ± 2.26 |
| <sup>b</sup>SM-D<sub>R,S</sub> | 9.56 ± 1.63 | 106 ± 7.19 |
| <sup>b</sup>SM-D<sub>S,R</sub> | 49.1 ± 11.4 | 22.1 ± 0.856 |
| <sup>c</sup>PF<sub>S,S</sub>-D<sub>R,S</sub> | 831 ± 99.5 | 282 ± 24 |
| <sup>c</sup>PF<sub>S,S</sub>-D<sub>S,R</sub> | 87.8 ± 9.81 | 1.85 ± 0.137 |

## Supporting Information

### Resolution of PF592,379 diastereomers and chemical synthesis of SM-C*_cis_* and SM-C*_trans_*

**Scheme S1.**
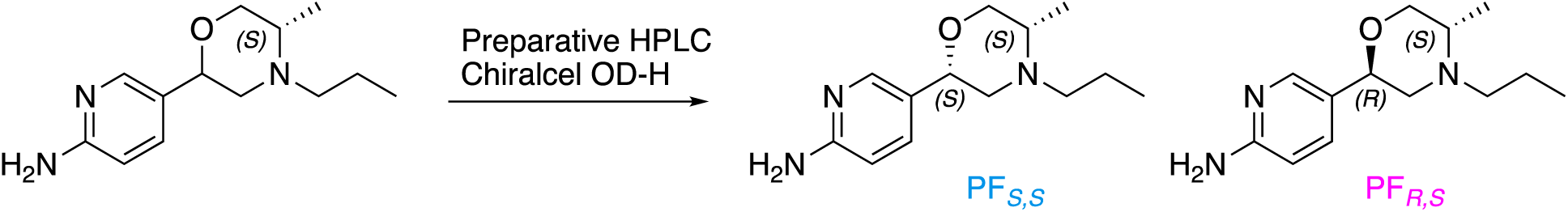
Resolution of (2*S*,5*S*)- and (2*R*,5*S*)-PF592,379 diastereoisomers.

The diasteromeric mixture (30.5 mg; 2.31 mmol), synthesized as previously reported ([10]; International Patent: WO2006/082511), was resolved by preparative chiral HPLC: Chiralcel OD-H (21 mm x 250 mm x 5 μm); mobile phase: isocratic 10% iPrOH in n-hexane + 0.1% DEA; temperature: 25 °C; flow rate: 15−18 mL/min; injection volume: 3 mL (∼5-10 mg/mL sample concentration); detection at λ 254 nm and 280 nm with the support of ELS detector. **PF*_S,S_*** eluted first (9.80 mg) and **PF*_R,S_*** eluted second (7.2 mg). Their spectroscopic data matched with the published literature and patents ([8, 10]; International Patent: WO2006/082511). Analytical chiral HPLC analysis performed using the Chiralcel OD-H analytical column (4.5 mm x 250 mm x 5 μm particle size); mobile phase: isocratic 10% iPrOH in n-hexane + 0.1% DEA; flow rate: 1 mL/min; injection volume: 20 μL; and sample concentration: ∼1 mg/mL. Multiple DAD λ absorbance signals were measured in the range of 210−280 nm. **PF*_S,S_*** *t*_R_ 11.795 min, purity > 99%, de > 99%; **PF*_R,S_*** *t*_R_ 22.343 min, purity > 99%, de > 99%.

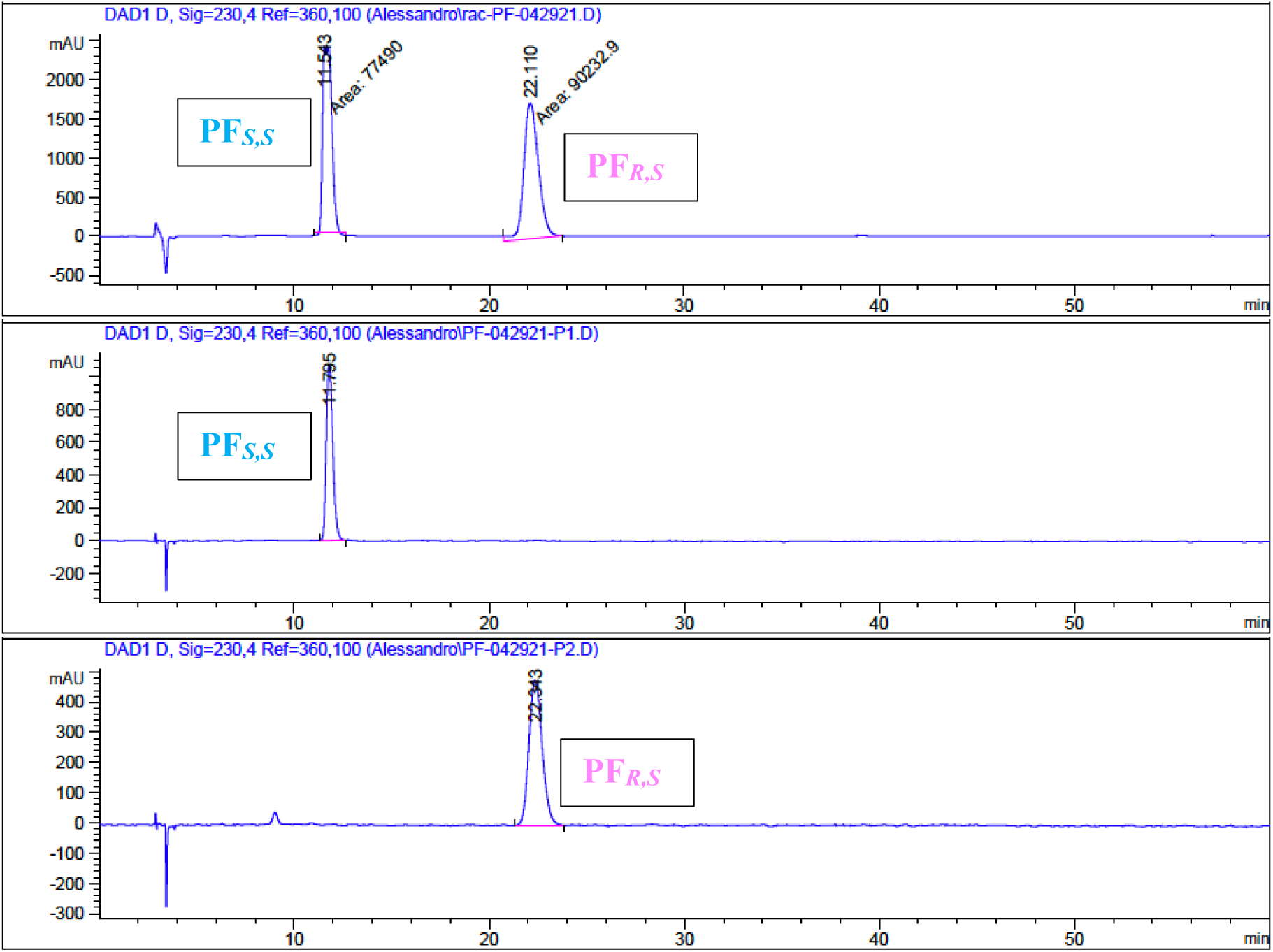

**Scheme S2.**
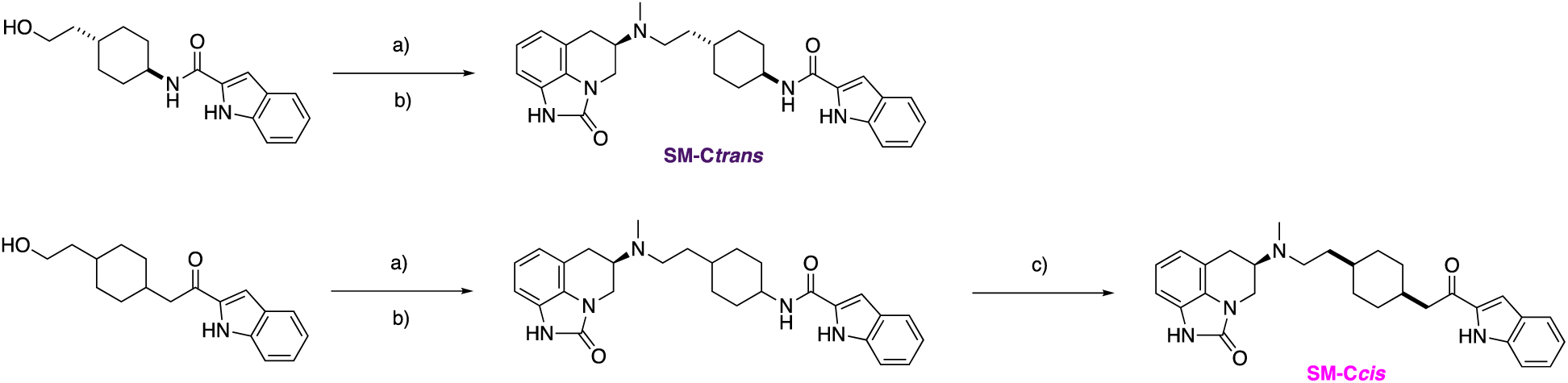
a) Dess-Martin periodinane (DMP), DCM, from 0 °C to RT; b) Sumanirole, cat. AcOH, NaBH(OAc)_3_ (STAB), 1,2-dichloroethane (DCE), RT; c) Preparative Chiral HPLC (Chiralpak-ADH).

### *Trans*-(*R*)-*N*-(4-(2-(methyl(2-oxo-1,2,5,6-tetrahydro-4*H*-imidazo[4,5,1-*ij*]quinolin-5-yl)amino)ethyl)cyclohexyl)-1*H*-indole-2-carboxamide (SM-C*_trans_*)

*Trans*-*N*-4-(2-hydroxyethyl)cyclohexyl)-1*H*-indole-2-carboxamide (17 mg; 0.06 mmol) was dissolved in 5 mL of DCM and cooled at 0 °C, and subsequently DMP (31 mg; 0.07 mmol) was added portion-wise to the reaction mixture. The reaction was removed from the ice bath and allowed to stir for 25 min at RT after which saturated aq. NaHCO_3_ was added to the reaction mixture. The aq. and organic layers were separated, the aq. layer extracted with DCM and the combined organic layers were dried with Na_2_SO_4_ and concentrated. The crude residue was used without further purification. The crude material was redissolved in 10 mL of DCE, followed by the addition of Sumanirole (10 mg, 0.05 mmol) and 3−4 drops of acetic acid. The reaction mixture was allowed to stir for 25 min, followed by the addition of STAB (16 mg, 0.07 mmol). The reaction was subsequently stirred overnight. The solvent was removed under reduced pressure, and the crude material was purified via flash chromatography with a gradient solvent system ramping from 0 to 20% DMA (MeOH in DCM + 1% aq. NH_4_OH) to yield the desired product (13 mg, 0.03 mmol, 56% yield). Analytical chiral HPLC analysis performed using the Chiralpak AD-H analytical column (4.5 mm x 250 mm x 5 μm particle size); mobile phase: isocratic 50% iPrOH in n-hexane; flow rate: 1 mL/min; injection volume: 20 μL; and sample concentration: ∼1 mg/mL. Multiple DAD λ absorbance signals were measured in the range of 210−280 nm. **SM-C*_trans_*** *t*_R_ 35.328 min, purity > 99%, de > 99%. ^1^H-NMR (400 MHz, CDCl_3_) δ 9.32 (br s, 1H), 8.55 (br s, 1H), 7.64 (d, *J* = 7.4 Hz, 1H), 7.44 – 7.39 (m, 1H), 7.27 - 7.31 (m, 1H), 7.16 – 7.08 (m, 1H), 6.96 - 6.78 (m, 4H), 6.00 (br s, 1H), 4.18 (d, *J* = 12.1 Hz, 1H), 4.00 – 3.86 (m, 1H), 3.57 (t, *J* = 11.7 Hz, 1H), 3.26 – 3.17 (m, 1H), 2.96 (br s, 2H), 2.70 – 2.53 (m, 2H), 2.39 (s, 3H), 2.11 (d, *J* = 12.5 Hz, 2H), 1.81 (d, *J* = 13.3 Hz, 2H), 1.72 – 1.54 (m, 3H), 1.30 - 1.10 (m, 4H). HRMS-ESI (MS/MS) C_28_H_33_N_5_O_2_ + H^+^ calculated 472.27070, found 472.27012.

### *Cis*-(*R*)-*N*-(4-(2-(methyl(2-oxo-1,2,5,6-tetrahydro-4*H*-imidazo[4,5,1-*ij*]quinolin-5-yl)amino)ethyl)cyclohexyl)-1*H*-indole-2-carboxamide (SM-C*_cis_*)

The desired product was synthesized following the same procedure described for SM-C*_trans_*, starting from *N*-4-(2-hydroxyethyl)cyclohexyl)-1*H*-indole-2-carboxamide (17 mg; 0.06 mmol). The desired product was obtained as *cis* and *trans* diastereomeric mixture (15 mg, 0.03 mmol, 65% yield; dr *cis*:*trans* 25:75). The *cis* diastereoisomer was isolated by preparative chiral HPLC: Chiralpak AD-H (21 mm x 250 mm x 5 μm); mobile phase: isocratic 50% iPrOH in n-hexane; temperature: 25 °C; flow rate: 15−18 mL/min; injection volume: 3 mL (∼5-10 mg/mL sample concentration); detection at λ 254 nm and 280 nm with the support of ELS detector. SM-C*_cis_* eluted first (3.5 mg) and SM-C*_trans_* eluted second and matched with the same *trans* reference standard described above. Analytical chiral HPLC analysis performed using the Chiralpak AD-H analytical column (4.5 mm x 250 mm x 5 μm particle size); mobile phase: isocratic 50% iPrOH in n-hexane; flow rate: 1 mL/min; injection volume: 20 μL; and sample concentration: ∼1 mg/mL. Multiple DAD λ absorbance signals were measured in the range of 210−280 nm. SM-C*_cis_ t*_R_ 24.317 min, purity > 99%, de > 99%. ^1^H-NMR (400 MHz, CDCl_3_) δ 9.32 (s, 1H), 8.44 (s, 1H), 7.68 – 7.61 (m, 1H), 7.47 – 7.43 (m, 1H), 7.32 – 7.28 (m, 1H), 7.18 – 7.09 (m, 1H), 7.00 – 6.92 (m, 1H), 6.91 – 6.82 (m, 3H), 6.25 (br s, 1H), 4.23 – 4.14 (m, 2H), 3.62 – 3.52 (m, 1H), 3.23 (br s, 1H), 2.99 – 2.91 (m, 2H), 2.71 – 2.56 (m, 2H), 2.41 (s, 3H), 1.84 - 1.24 (m, 11H). HRMS-ESI (MS/MS) C_28_H_33_N_5_O_2_ + H^+^ calculated 472.27070, found 472.27011.

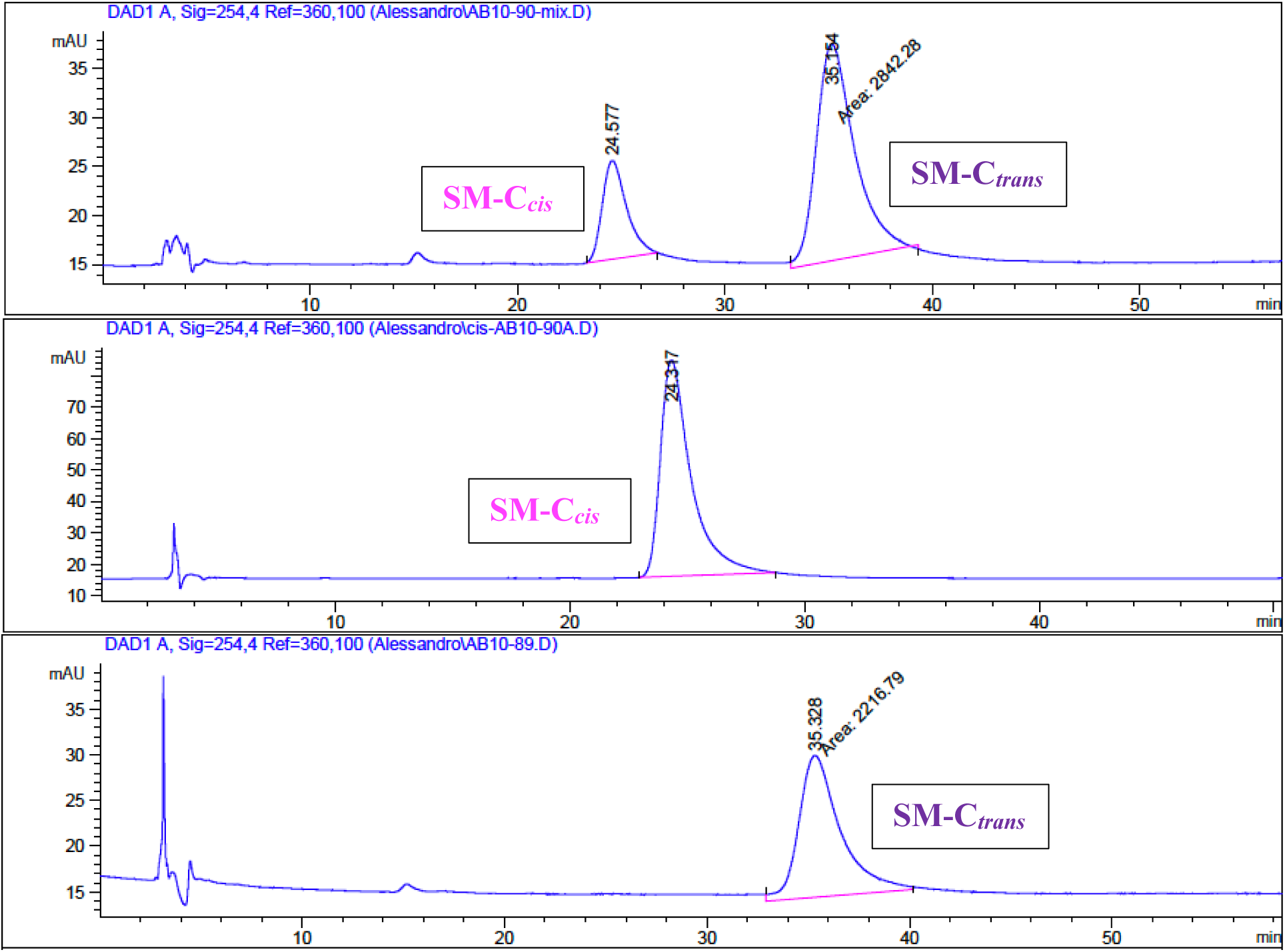

